# The coagulation factor IX (F9) loss of function prevents the cell cycle arrest induced by CDK4/6 inhibitors treatment

**DOI:** 10.1101/2021.11.30.470520

**Authors:** Paula Carpintero-Fernández, Michela Borghesan, Olga Eleftheriadou, Juan Antonio Fafián-Labora, Tom P. Mitchell, Tom D. Nightingale, María D. Mayán, Ana O’Loghlen

## Abstract

During this last decade the development of pro-senescence therapies has become an attractive strategy as cellular senescence acts as a barrier against tumour progression. In this context, CDK4/6 inhibitors induce senescence and have showed efficacy in reducing tumour growth in breast cancer patients. However, even though cancer cells are arrested after CDK4/6 inhibitor treatment, genes regulating senescence in this context are still unknown limiting their anti-tumour activity. Here, using a functional genome wide CRISPR/Cas9 genetic screen we found several genes that synergistically participate in the proliferation arrest induced by the CDK4/6 inhibitor, Palbociclib. We find that downregulation of the coagulation factor IX (*F9*) using sgRNA and shRNA prevents the cell cycle arrest and senescent-like phenotype induced in MCF7 breast tumour cells upon Palbociclib treatment. These results were confirmed using another breast cancer cell line and with an alternative CDK4/6 inhibitor, Abemaciclib, and further tested in a panel of 22 cancer cells. While *F9* knockout reduces senescence, treatment with a recombinant F9 protein was sufficient to induce a cell cycle arrest and senescence-like state in MCF7 tumour cells. Besides, endogenous F9 is upregulated in different human primary cells cultures undergoing senescence. Importantly, bioinformatics analysis of cancer datasets suggest a role for F9 in human tumours. Altogether, these data collectively propose key genes involved in CDK4/6 inhibitors response that will be useful to design new therapeutic strategies in personalized medicine in order to increase their efficiency, stratify patients and avoid drug resistance.

## INTRODUCTION

A key characteristic of cancer cells is the deregulation of cell cycle checkpoint proteins such as the cyclin-dependent kinases (CDKs) CDK4 and CDK6 leading to uncontrolled cell proliferation. Molecular changes at CDKs level have been reported in various cancer types making them an attractive potential target for new treatments^1,2^. CDK inhibitors, in particular CDK4/6 inhibitors (Abemaciclib, Palbociclib and Ribociclib) induce a cell-cycle arrest and subsequently activate senescence in many human cancer cell lines^3,4,5,6,7,8^. Also, these inhibitors have been recently reported to promote anti-tumour immunity^9^, reduce NADPH and glutathione levels^10^ and stimulate tumour antigen presentation^11^. Although the three CDK4/6 inhibitors reached phase III clinical trials, Palbociclib progressed towards the clinic, receiving accelerated approval from the Food and Drug Administration (FDA) in February 2015^12^ for estrogen-receptor positive (ER^+^)/HER2 negative (HER2^-^) breast cancer subtypes. Abemaciclib was FDA approved in 2017 to be used either alone^13^ or in combination with fulvestrant for women with ER^+^/HER2^-^ advanced or metastatic breast cancer with disease progression following endocrine therapy^14,15^.

Palbociclib (PD-0332991), a second-generation CDK4/6 inhibitor, has shown effectiveness specially in advanced ER^+^/HER2^-^ breast cancer^16^, improving the patient progression-free survival from 18 to 27 months^17^. Palbociblib showed beneficial effects compared to hormone therapy, letrozole (an aromatase inhibitor) or fulvestrant (an ER^+^ antagonist) when using alone^16,18^. Importantly, there are several clinical trials undergoing employing Palbociclib in a variety of other cancer types such as squamous cell lung cancer (NCT02785939), pancreatic neuroendocrine tumours (NCT02806648) and oligodendroglioma and oligoastrocytoma (NCT02530320). However, not all patients respond to Palbociclib treatment suggesting that mechanisms that drive resistance or prevent the expected response exist, highlighting the importance to develop more personalized cancer therapies^15^. Mechanistically, Palbociclib inhibits the phosphorylation of retinoblastoma (RB1), stabilizing the RB1-E2F inhibitory complex and preventing the activity of E2F transcription factor family that regulates cell cycle progression and apoptosis^19^. In fact, Palbociclib induces cellular senescence by inducing a G1 arrest and inhibits growth of tumour xenographs *in vivo*^20,21^. Furthermore, loss of RB1 function is an established mechanism of primary resistance to CDK4/6 inhibitors *in vitro*^15,19,22^. However, further research is required to identify new biomarkers of resistance to these inhibitors.

Cellular senescence is defined as a state in which cells lose their proliferative capacity despite them being metabolically active. Senescent cells participate in a wide range of biological processes playing both beneficial or detrimental effects for the organism^23,24^. As part of the senescence program, senescent cells differ from dividing cells in terms of gene expression, chromatin structure and metabolism^24,25^ which makes them susceptible to certain drugs that do not affect their proliferating counterparts^26^. Senescent cells also comprise a complex of pro-inflammatory response proteins known as senescence-associated secretory phenotype (SASP)^27^. The SASP is characterised by the secretion of cytokines, enzymes, and chemokines that cause inflammation and is pivotal for the clearance of senescent cells by phagocytosis^28^. However, the role the SASP plays during cancer is still under debate^24,25^. While some studies show its beneficial effects as a tumour suppressor mechanism, others have demonstrated the SASP promotes tumorigenesis^29,30^. Although most primary cell types follow the senescence program and have beneficial effects for the microenvironment^23^, cancer cells tend to overcome senescence resulting in uncontrolled cellular proliferation and tumorigenesis^24,25^. Thus, the molecular mechanisms by which CDK4/6 inhibitors induce senescence in cancer cells and the genes involved in conferring drug resistance are unknown. This lack of knowledge prevents the stratification of patients prior to Palbociclib treatment or to develop therapeutic strategies to avoid drug resistance in order to increase the progression free survival of cancer patients.

In this study, using a human genome-wide CRISPR/Cas9 library we identified genes whose loss of function prevent the proliferative arrest induced by Palbociclib in MCF7 breast cancer cell line. Validation of the CRISPR/Cas9 screen using four independent sgRNA and two individual shRNA confirmed that among the identified genes, the coagulation factor IX (*F9*) participates in the cell cycle arrest induced by Palbociclib. These results were established using Abemaciclib where we saw that downregulation of *F9* also prevented the induction of senescence. Meanwhile, treatment of the breast cancer cell line MCF7 with recombinant F9 induced a senescence-like proliferative arrest. Our results demonstrate that *F9* mRNA is endogenously upregulated upon the acquisition of the senescent phenotype as shown during oncogene-induced senescence (OIS), DNA damage-induced senescence (DDIS) and therapy-induced senescence (TIS) in human primary fibroblasts. Furthermore, treatment of human primary endothelial cells with Palbociclib mimicked a senescence-like phenotype increasing the expression of F9. Finally, we screened a panel of 22 cancer cell lines for their response to different CDK4/6 inhibitors and we show that *F9* loss of function confers a partial resistance to the proliferative arrest induced by CDK4/6 inhibitors in other tumour types. Analyses of published datasets also suggest a role for F9 in carcinogenesis in different tumours in humans. Importantly, our results open new therapeutic opportunities with the potential to stratify patients for CDK4/6 inhibitors response prior to treatment.

## RESULTS

### A CRISPR/Cas9 genome-wide screen identifies genes regulating the proliferative arrest induced by Palbociclib treatment

In order to identify genes whose loss of function (LOF) overcome the proliferative arrest induced by Palbociclib (PD-0332991 or Palbo hereafter) we performed a CRISPR/Cas9 screen using the human genome-wide library GeCKO*v2*^31,32^. This library contains ∼ 123,441 unique sgRNA targeting 19,050 genes in the human genome with a coverage of 5-6 sgRNA per gene (**Figure 1A**). Firstly, to confirm that treatment with Palbo induces senescence we treated the ER^+^ breast cancer cell line MCF7 with increasing concentrations of Palbo (0.1, 0.2, 0.5 and 1μM) for 7 and 14 days and analysed a variety of markers characteristic of senescence. We confirmed that treatment with Palbo induced a stable cell cycle arrest by quantifying the number of cells staining positive for BrdU in addition to determining the relative cell number (**Figure S1A**). Furthermore, Palbo treatment induced an increased in lysosomal activity, a common characteristic of senescence by measuring β-Galactosidase activity (SA-β-Gal) (**Figure S1B**). Neither of the doses used induced apoptosis quantified by measuring the number of cells staining positive for AnnexinV (**Figure S1C**). We next determined additional markers of senescence by treating MCF7 with 200nM Palbo during 14 days and confirmed SA-β-Gal, in addition to an increase in the number of cells staining positive for p21^CIP^ by immunofluorescence (**Figure S1D-F**). Next, we infected MCF7 cells with a single-vector lentiviral construct (comprising sgRNA and human Cas9 in a single vector), lentiCRISPRv2, empty vector (**Figure S1G**) or containing the GeCKO pooled library and treated them with DMSO or 200nM Palbo for 14 days (**Figure 1A**). As proof of concept that the screen worked, MCF7 cells were plated at low density to determine their proliferative capacity where an advantage in proliferation can be observed upon the expression of the GeCKO library (**Figure 1B**). Enrichment of sgRNAs after two weeks Palbo treatment compared to day 0 was determined by genomic DNA extraction and deep sequencing as previously described^31,32^ (**Figure 1C**). Among all the sgRNA enriched after two weeks treatment (p<0.05) we selected: (i) single sgRNA enriched more than ≥ 2 log_2_ fold change RPKM between day 0 and 14 and, (ii) those sgRNA where we found more than ≥ 3 individual sgRNA per gene preventing the proliferative arrest. This gave us a list of 18 potential genes whose loss of function prevent the cell cycle arrest induced by Palbo (**Figure 1D**). We subjected these 18 genes to KEGG (Kyoto Encyclopedia of Genes and Genomes) pathway and STRING protein interaction analysis and strikingly we noted a number of genes associated with the blood coagulation pathway (**Figure 1E,1F**). Within these genes we found overrepresentation of the coagulation factor IX (*F9*) and Protein Z Vitamin K Dependent Plasma Glycoprotein (*PROZ*) (**Figure 1D, 1E**). Enrichment of the individual sgRNA within the GeCKO library belonging to the coagulation pathway (5 sgRNAs for *PROZ* and 6 sgRNA for *F9*) with ≥ 2 log_2_ fold change RPKM after two weeks Palbo treatment show a statistical difference (**Figure 1G**). Altogether, these data propose 18 candidate genes whose loss-of-function prevent the proliferation arrest induced by Palbo with an overrepresentation of two genes involved in the blood coagulation pathway.

**Figure 1.**
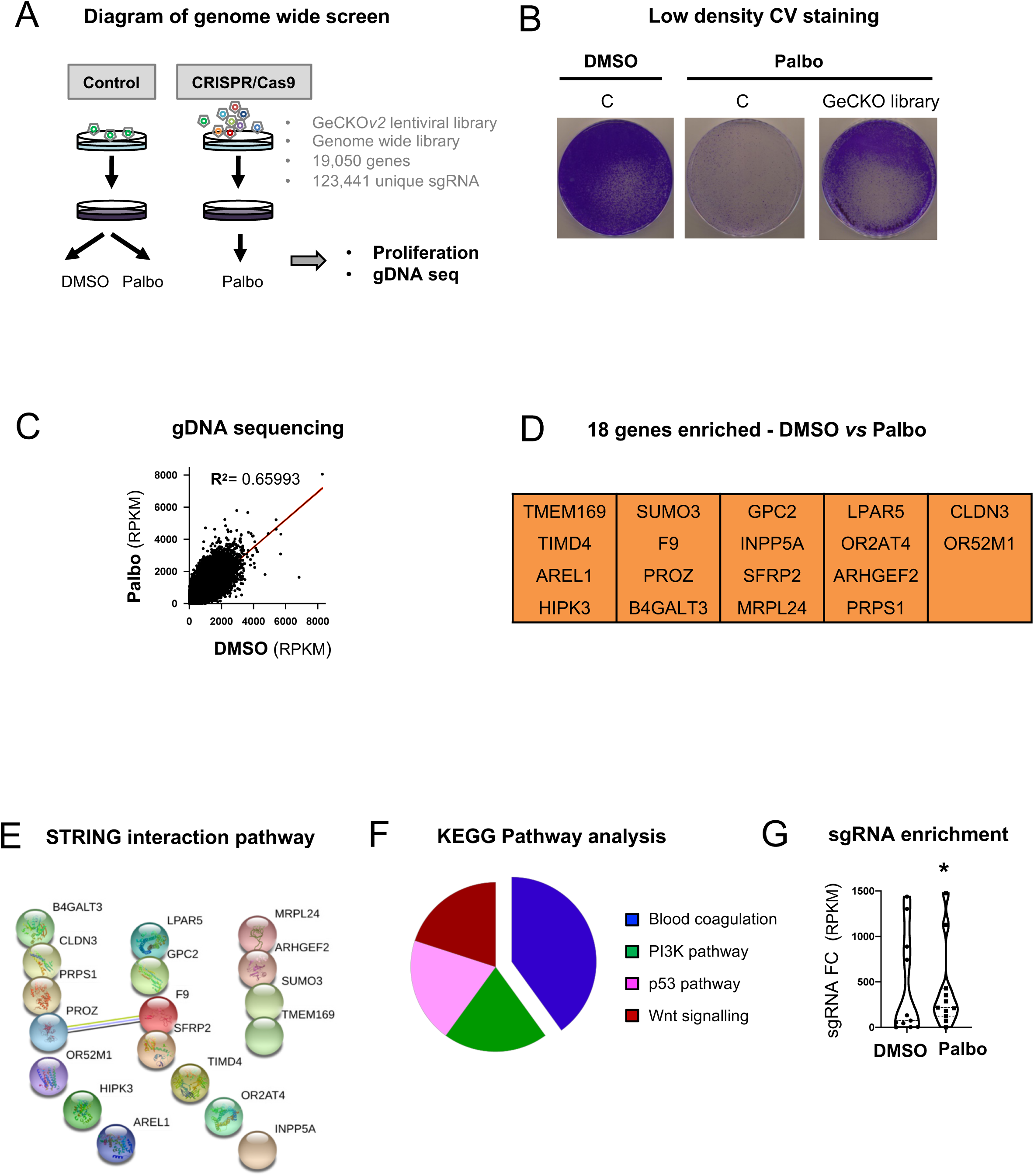
CRISPR/Cas9 screening identifies candidate genes implicated in Palbociclib cell cycle arrest. **(A)** Schematic representation of the proof-of-concept genome wide screen performed in MCF7 using the GeCKO*v2* pooled sgRNA library. Cells were infected with the library (CRISPR/Cas9) or the vector control (control), selected with puromycin and treated with 200nM of Palbociclib (Palbo) for 14 days. **(B)** MCF7 cells expressing either the empty vector (C) or the GeCKO library after 14 days of 200nM Palbo treatment were stained with crystal violet. A representative experiment of 2 independent experiments is shown. **(C)** Genomic DNA (gDNA) sequencing data showing the enrichment of sgRNA after two weeks of 200nM Palbo treatment. Data show a representative experiment from 2 independent experiments. Statistically significant (p<0.05) transformed RPKM is shown. **(D)** sgRNA targeting 18 different genes were found to be statistically significant following the selection criteria of: (i) >2 FC (fold change) differential expression between DMSO and day 14 Palbo treatment and, (ii) 3 or more sgRNA conferring a proliferative advantage. **(E)** STRING protein interaction and **(F)** Kyoto Encyclopedia of Genes and Genomes (KEGG) analysis for the 18 genes whose sgRNA were enriched after 14 days Palbo treatment in panel D. **(G)** Violin plot showing all individual sgRNA within the GECKO library related to the coagulation pathway (5 sgRNAs for *PROZ* and 6 sgRNA for *F9*) enriched after two weeks Palbo treatment (FC, fold change RPKM). Median for values is shown for all sgRNA from 2 independent experiments. One sample t and Wilcoxon test was performed. Related to **Figure S1**.

### *F9* and *PROZ* loss of function prevents the proliferation arrest induced by Palbo

It has been recently reported that chemotherapy-induced senescence potentiates blood clotting by modifying platelet function^33^. Among the genes identified in our screen we found overrepresentation of two factors participating in the blood coagulation pathway (**Figure 1D-G**), thus we decided to focus on these genes for further validation. For this we designed and cloned 4 additional sgRNA sequences different from those in the GeCKO library targeting *PROZ* (sgPROZ) and *F9* (sgF9). We also included an sgRNA targeting *RB1* (sgRB) as a positive control, as cancer cells lacking this gene generally do not respond to CDK4/6 inhibitors^16^. MCF7 cells were infected with the 4 sgRNAs targeting each candidate gene. To determine their proliferative capacity, MCF7 cells were treated with either DMSO or 200nM Palbo and cell numbers were determined on days 6, 12 and 20 after Palbo treatment (**Figure 2A**). The ability of sgRNAs targeting *F9*, *PROZ* and *RB1* to prevent the proliferation arrest induced by Palbo, was confirmed by proliferation curves at different days as shown in **Figure 2B**. In fact, sgF9 and sgPROZ bypassed the cell cycle halt induced by Palbo when compared with MCF7 control cells at day 20 (**Figure 2B**). The efficacy of the different sgRNAs was assessed at the mRNA level by qPCR (**Figure S2A**) and protein level for RB (**Figure S2B**). Furthermore, we established that the basal cell proliferation rate was not affected by the sgRNA expression in comparison with the control cells (**Figure 2C**) confirming that the bypass in proliferation was specific to sgF9 and sgPROZ upon Palbo treatment. We further established we could observe an increase in the mRNA expression levels of *F9* and *PROZ* in MCF7 cells after treating with Palbo for 20 days (**Figure 2D**). Next, as F9 can be secreted we determined whether we could detect it in the conditioned media upon in senescent cells treated with Palbo. As expected, we could confirm an increase in the amount of F9 released upon Palbo treatment by ELISA (**Figure 2E**), suggesting that F9 could be part of the SASP.

**Figure 2.**
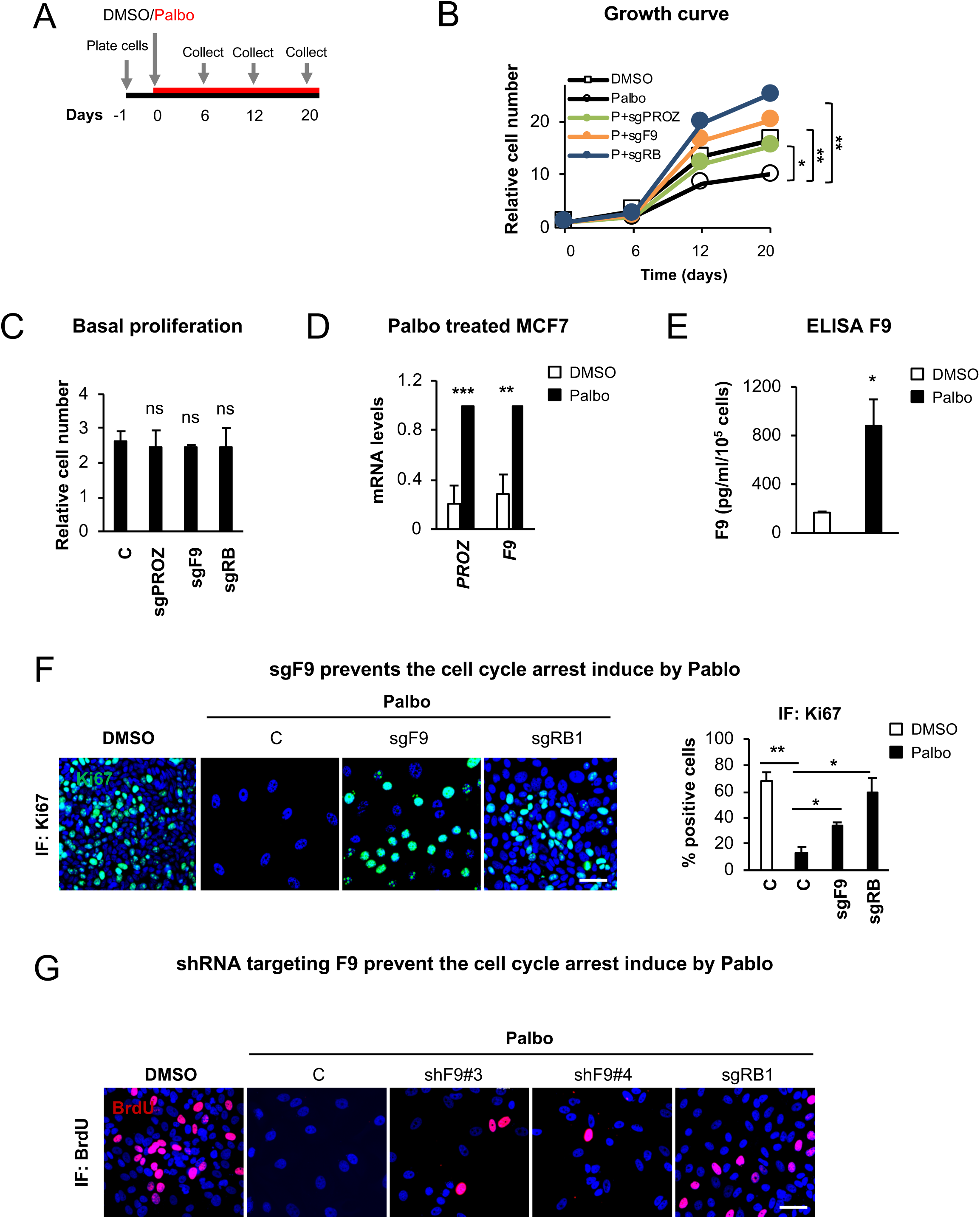
CRISPR/Cas9 screen validation identifies *F9* as a regulator of the proliferation arrest induced by Palbo. **(A)** Overview of the experimental set-up followed to validate the identified sgRNA. Briefly, after plating, MCF7 cells were treated with 200nM Palbo and samples were collected to determine cell number at days 6, 12 and 20 after Palbo treatment. **(B)** MFC7 cells expressing the indicated sgRNAS were treated with 200 nM Palbo for 20 days and collected at different timepoints (0, 6, 12, 20 days) to asses proliferation. Proliferation curves show that MCF7 expressing sgF9 (orange line) and sgPROZ (green line) prevented a stable cell cycle arrest compared to Palbo (P) treated cells (black line - circles). sgRB MCF7 cells treated with Palbo (blue line) were used as a positive control. The data represent the mean of 3-5 independent experiments. Student’s t-test analysis at day 20 was performed compared to the Palbo treated sample. **(C)** Basal proliferation rate was determined by quantifying nuclei count after different sgRNA infections at 20 days of cell culture. Data show the mean ± SEM of 2-3 independent experiments. Two-tailed students t-test compared to the C sample was performed. **(D)** MCF7 treated with Palbo for 20 days induce an upregulation of *F9* and *PROZ* mRNA levels as shown by qPCR analysis. Data represent the mean ± SEM of 5 independent experiments. Two-tailed t-test analysis was performed. **(E)** ELISA for F9 protein levels secreted by MCF7 cells upon DMSO or Palbo treatment for 20 days. Data represent the mean ± SEM of 4 independent experiments. Two-tailed t-test was performed. **(F)** Representative images and quantification showing that sgF9 prevents the proliferation arrest induced by Palbo by displaying an increase in the percentage of cells staining positive for Ki67 (green). Data show the mean ± SEM of 3 independent experiments. Two-tailed t-test analysis comparing to Palbo sample was performed. Scale bar: 50μm. **(G)** Representative images for BrdU staining of MCF7 cells infected with a construct expressing two individual shRNA targeting F9 (shF9#3 and shF9#4). sgRB is used as a positive control. Scale bar: 50μm. See also **Figure S2**.

We next wanted to confirm that the proliferation arrest upon Palbo treatment was stable, a characteristic of senescence, and that both sgPROZ and sgF9 were implicated in bypassing this arrest, thus not due to spontaneous hyperproliferation. For this, we treated MCF7 cells for 6 days with Palbo, washed the plates and cultured them further in the absence of this drug until day 20 (**Figure S2C**). As shown in **Figure S2D** treatment of MCF7 cells with Palbo resulted in a stable inhibition of proliferation even in the absence of the drug while the expression of both sgF9 and sgPROZ prevented this proliferation arrest as shown by low density MCF7 plating and crystal violet staining (**Figure S2D**).

### *F9* loss of function prevents the senescence-like phenotype induced by Palbo

Next, we decided to focus on *F9* knockout as its proliferation bypass is stronger than sgPROZ (**Figure 2B**). We thus determined whether this proliferative advantage was maintained using different concentrations of Palbo. For this, we treated MCF7 cells with 200nM and 500nM Palbo for 20 days and the relative cell number was determining at day 20 (**Figure S2E**). The knockout efficiency of sgF9 and sgRB were tested after day 20 Palbo treatment to ensure the plasmid expression was not lost (**Figure S2F).** Additional methods were used to determine the bypass mediated by sgF9 such as quantifying the percentage of cells staining positive for Ki67 by IF (**Figure 2F**) and low density cell plating and crystal violet staining (**Figure S2G**). We next tested whether sgF9 could be inducing an increase in the migration capacity of MCF7 cells, thus promoting tumorigenesis. However, migration assays show a decrease in the relative number of MCF7 cells migrating upon 500nM Palbo treatment for 20 days which is not prevented by the loss of F9 (**Figure S2H**). MDA-MD- 468 cells were used as a positive control due to their high migration capacity.

To further confirm the specificity of sgF9 in bypassing Palbo growth inhibition and to exclude the potential implication of off-target effects derived from using CRISPR/Cas9 technique, we infected MCF7 cells with two independent viral constructs carrying an shRNA targeting *F9* (shF9#3 and shF9#4) and treated MCF7 cells with 500nM Palbo for 20 days. Both shF9 constructs recapitulated the effects observed with the sgRNA targeting *F9* measured by quantifying the number of cells staining positive for BrdU (**Figure 2G**). *F9* downregulation using both shF9 constructs was confirmed at the mRNA level by qPCR (**Figure S2I**). Altogether, these data show that *F9* loss of function using both genome editing and RNAi interference techniques overcomes the cell cycle arrest induced by Palbo treatment in MCF7 cells.

### F9 is endogenously upregulated during senescence

Next, we set to determine the importance of F9 expression during senescence; thus, we evaluated whether *sgF9* prevented the induction of other features of senescence induced by Palbo such as the SASP. We can observe that *sgF9* downregulates several SASP mRNA transcripts upregulated by the treatment with 500nM Palbo such as *MMP9*, *MMP3*, *IL1B* and *IL6* while having no effect on *CCL20*, *IL1A* or *IL8* (**Figure 3A**). Importantly, sgRB which is known to prevent the activation of senescence averted the endogenous upregulation of *F9* mRNA levels (**Figure 3B**), suggesting that *F9* mRNA expression is dependent on RB1. As oncogene-induced senescence (OIS) is a potent tumour suppressor mechanism both *in vitro* and *in vivo*, we next took advantage of human primary fibroblasts (HFFF2) expressing an endoplasmic reticulum (ER):H-RAS^G12V^ fusion protein (iRAS). Upon treatment with 200nM 4-hydroxytamoxifen (4OHT) for 6 days, senescence is progressively established^8,34^. By quantifying the mRNA levels of *F9*, we can observe a consistent upregulation of endogenous *F9* during OIS (**Figure 3C, left panel**). In addition, we confirmed that the upregulation of *F9* transcript was accompanied by increased levels in SA-β-gal activity upon iRAS induction compared to the vector control (**Figure S3A**). We also observed an upregulation of endogenous *F9* mRNA levels in HFFF2 during DNA-damage-induced senescence (DDIS) induced by treating HFFF2 with 50μM etoposide for 2 days and collecting for RNA at day 7 (**Figure 3C, middle panel**). Moreover, treatment of HFFF2 fibroblasts with 1μM Palbo for 7 days, mimicking therapy-induced senescence (TIS), also triggered endogenous upregulation of *F9* mRNA (**Figure 3C, right panel**). Confirmation of the induction of senescence was further demonstrated by showing a reduction in cell proliferation measured by quantifying cell number (**Figure S3B**) and an increase in the number of SA-β-gal positive cells (**Figure S3C**) as previously^8,34,35^.

**Figure 3.**
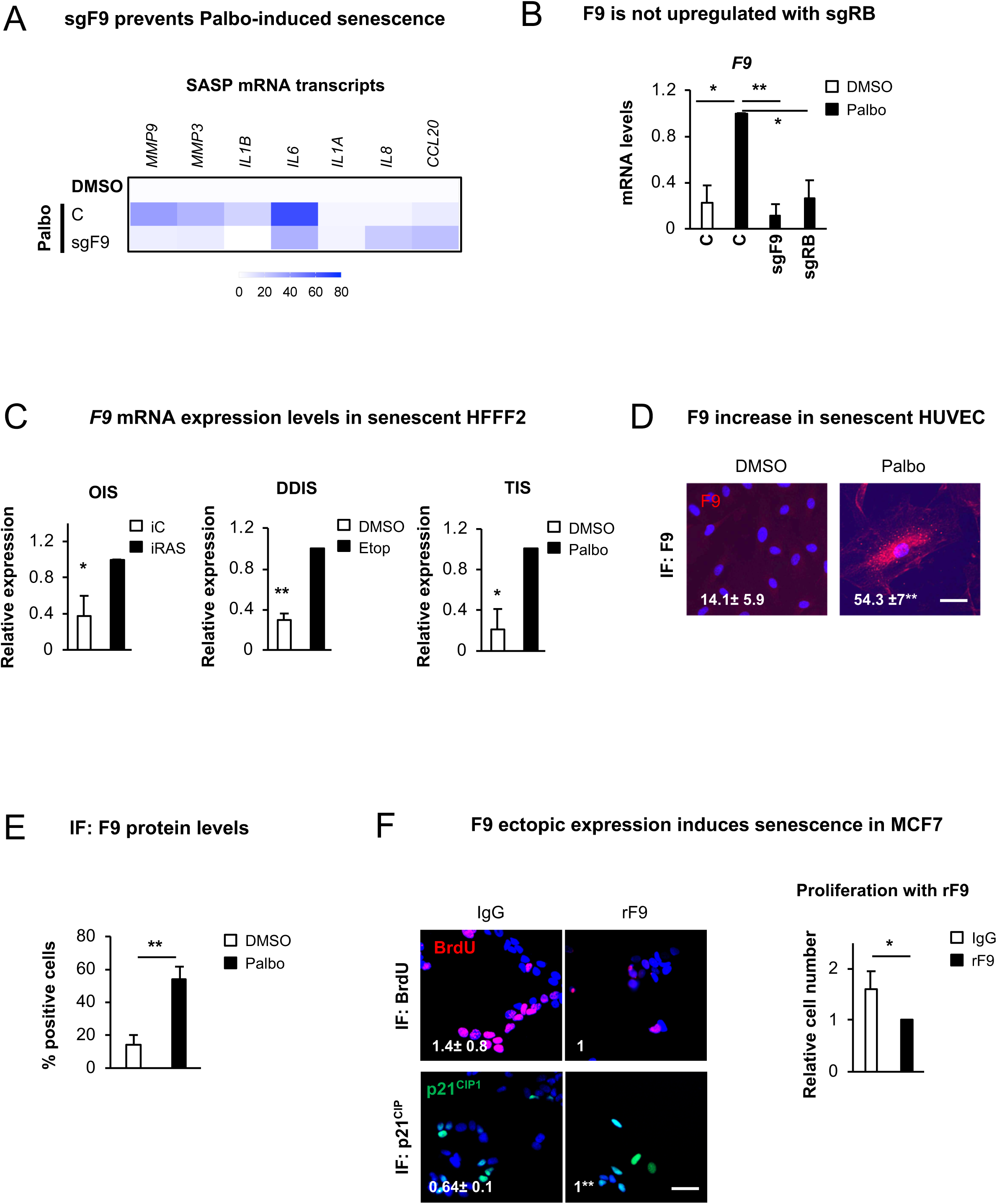
F9 induces senescence and it is endogenously upregulated during senescence. **(A)** Heat map of SASP mRNA levels in MCF7 cells control or expressing sgF9 after 20 days treatment with Palbo. The mean of 2-7 independent biological replicates is shown. **(B)** *F9* mRNA upregulation by Palbo is prevented when RB is not present (sgRB). Data show the mean ± SEM of 4 independent experiments. One-WAY ANOVA with Dunnett’s multiple comparison to Palbo sample was performed. **(C)** mRNA levels of endogenous *F9* mRNA levels in HFFF2 (human primary fibroblasts) upon the induction of senescence. OIS (Oncogene-induced senescence) was induced in HFFF2 expressing ER:H-RAS^G12V^ (iRAS) by adding 200nM 4OHT for 6 days (left panel); 50μM etoposide was added for 2 days and washed out until day 7 to induce DDIS (DNA-damage senescence) (middle panel); TIS (Therapy-induced senescence) was mimicked by treating with 1μM of Palbo for 7 days (right panel). Data represent the mean ± SEM of 2-3 independent experiments. Two-tailed student’s t-test analysis was performed. **(D)** Representative immunofluorescence images and **(E)** IF quantification showing the expression of endogenous F9 (red) in HUVEC (human umbilical vein endothelial cells) upon 7 days treatment with 500nM Palbo. Scale bar: 50μm. The data represent the mean ± SEM of 3- 4 independent experiments. Two-tailed Student’s t-test was used to calculate statistical significance. **(F) Left panel,** Representative images for BrdU (red) and p21^CIP1^ (green) in MCF7 treated twice with 10μg/mL of recombinant F9 (rF9) for 6 days. Scale bar: 50 μm. 3 independent experiments were performed. **Right panel,** The graph shows a reduction in the number of cells at the end of the experiment. Mean ± SD of 3 independent experiments is shown. Two-tailed Student’s t-test was used to determine statistical significance. See also **Figure S3**.

Uncontrolled coagulation contributes to the pathophysiology of several chronic inflammatory diseases. In these conditions senescent cells are often observed and participate in the generation of inflammation^36^. Besides, endothelial cell activation during disease depends on a broad range of inflammatory mediators released by platelets^37^. While the role of platelets in haemostasis and wound repair is well known^38^ the participation of senescent cells in haemostasis was not reported until recently^33^. On the basis of these observations, we wanted to evaluate if the induction of senescence in human umbilical vein endothelial cells (HUVEC) using Palbo increases the expression levels of F9. Consistent with the data obtained for MCF7 and HFFF2, HUVEC cells treated with 500nM Palbo for 7 days showed an increase in the percentage of cells staining positive for F9 as shown by IF and its relative quantification (**Figure 3D, 3E**) and a reduced proliferation quantified by determining the percentage of cells incorporating BrdU (**Figure S3D, S3E**). In parallel, we observed an increase in the protein levels of the cell-cycle inhibitor p21^CIP1^ (**Figure S3D**) and in the quantification of the number of cells incorporating BrdU (**Figure S3E**).

Finally, we tested if ectopic administration of F9 using recombinant F9 (rF9) protein, induced senescence in MCF7 cells. The results obtained demonstrate that the administration of 10μg/mL rF9 twice for 6 days induced a senescence-like phenotype in MCF7 cells shown by a reduction in the number of cells staining positive for BrdU concomitant with an increase in number of cells staining positive for p21^CIP^ (**Figure 3F**). We further confirmed the proliferation arrest quantifying the relative cell number upon rF9 treatment (**Figure 3F**). Altogether, these data highlight the implication of F9 during senescence induced by a variety of triggers in different primary cell cultures.

### *F9* regulates senescence in MCF7 cells treated with Abema and in T47D

Next, we wanted to determine whether the proliferation bypass by *F9* loss of function was due to the specific inhibition of CDK4/6 and not due to off target effects induced by Palbo. For this determined the ability of MCF7 cells to respond to increasing concentrations of other CDK4/6 inhibitors (Abemaciclib or Abema and Ribociclib or Ribo) (**Figure 4A**). We used increasing concentrations of Abema and Ribo (0.25, 1 and 5μM), and included Palbo as a positive control (**Figure 4A, 4B**). We could indeed observe a dose-dependent cell cycle arrest upon treatment with other CDK4/6 inhibitors (Abema in particular) (**Figure 4B**) which was maintained after 14 days of continuous treatment (**Figure S4A, S4B)**. We next confirmed the induction of a stable cell cycle arrest characteristic of senescence by treating MCF7 cells for 6 days with Abema, Palbo or Ribo, withdrawing the inhibitors and leaving for 14 further days after the drug removal (**Figure S4C**). Our data confirm that both Palbo and Abema induce a stable cell cycle arrest or senescence even after drug withdrawal which was not maintained when the cells were treated with Ribo (**Figure S4C**). In line with these results, we could observe by qPCR analysis upregulation of *F9* mRNA levels when MCF7 cells were treated with Palbo and Abema but not with Ribo (**Figure 4C**). To further confirm an implication for *F9* in overcoming the proliferative arrest induced by CDK4/6 inhibitors, we treated MCF7 cells infected with sgF9 for 20 days with 1μM Abemaciclib. Consistent with our previous results, sgF9 partially prevented the cell cycle arrest induced by Abema as shown by an increase in the number of cells stained by crystal violet (**Figure 4D**). Next, we determined the implication of sgF9 using another ER^+^ (T47D) and the triple negative (MDA-MB-468) breast cancer cell lines. sgF9 also prevented the cell cycle arrest induced by Palbo in T47D cells as shown by determining proliferation by colony formation assays and crystal violet staining (**Figure 4E**) and by counting relative cell numbers (**Figure 4F**). The knockout efficiency for *RB* and *F9* was determined at the protein level for RB (**Figure S4D**) and RNA levels for *RB* and *F9* (**Figure S4E)**. However, MDA-MB-468 cells did not respond to Palbo treatment as previously reported^20^ (**Figure 4E**) in spite of *F9* and *RB* being downregulated (**Figure S4F**) by their respective sgRNAs. The induction of senescence by Palbo treatment was confirmed in T47D also by determining SA-β-Gal activity and quantification (**Figure 4G**). The secretion of F9 protein to the conditioned media was also confirmed by ELISA in T47D cells upon 1μM Palbo treatment for 20 days (**Figure 4H**). The implication of *F9* loss of function in T47D cell lines was also validated using two independent shF9#3 and shF9#4 by crystal violet staining (**Figure S4G**) and relative cell counting (**Figure S4H**). Confirmation of the knockdown efficiency was determined by measuring the levels of *F9* upon shF9 infection in T47D cells (**Figure S4I**).

**Figure 4.**
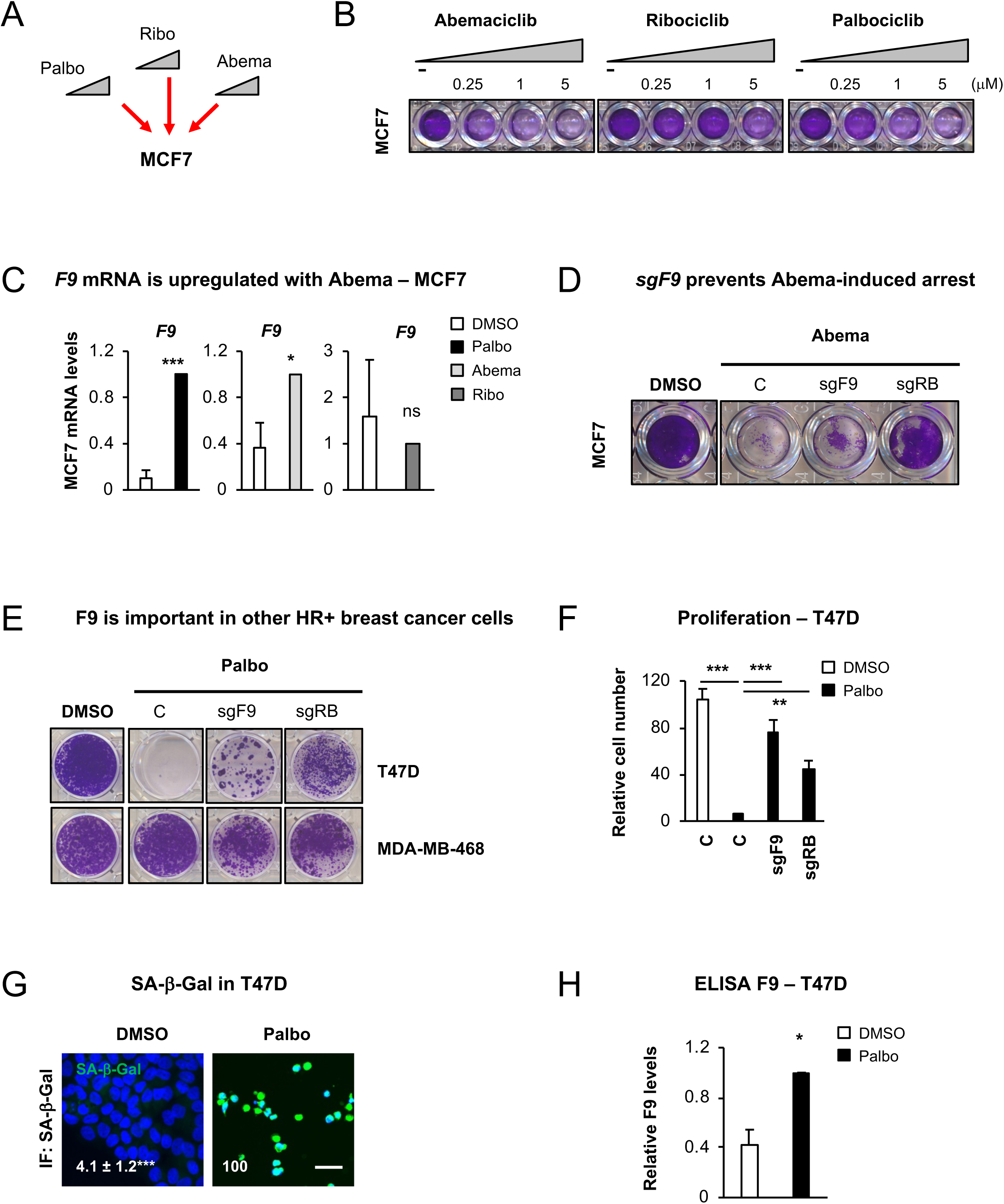
CDK4/6 inhibitors induce a cell cycle arrest in different tumour types and is dependent on *F9* in T47D cells. **(A)** Schematic representation of the treatment of MCF7 cells with 3 different CDK4/6 inhibitors: Palbociclib (Palbo), Ribociclib (Ribo) and Abemaciclib (Abema). **(B)** Colony formation assay stained with crystal violet for MCF7 treated with increasing concentrations of different CDK4/6 inhibitors for 10 days. A representative experiment is shown. **(C)** qPCR data show an upregulation of endogenous *F9* mRNA levels upon treatment with 500nM Palbo, 500nM Abema or 500nM Ribo. Data show the mean ± SEM of 3 independent experiments. Two-tailed t-test analysis is shown for statistical significance. **(D)** Crystal violet staining for colony formation in MCF7 cells expressing sgF9 or sgRB treated with 1μM Abema for 20 days. Representative experiment is shown. **(E)** Crystal violet staining shows the effect on proliferation in T47D (ER^+^ breast cancer cells) and MDA-MB-468 (triple negative breast cancer cell line) expressing sgF9 or sgRB and treated with Palbo. Representative staining of 4 independent experiments is shown. **(F)** Relative cell count for T47D cells expressing either sgF9 or sgRB. Data show the mean ± SEM of 7 independent experiments. One Way ANOVA with Dunnett’s multiple comparisons to Palbo C sample was performed. **(G)** Representative pictures and quantification for number of cells presenting SA-β-Gal activity in T47D treated with 1μM Palbo for 20 days. Two-tailed student’s t-test was performed. Scale bar: 50 μM. **(H)** ELISA for human F9 released to the conditioned media in T47D cell treated with 1μM Palbo for 20 days. Two-tailed student’s t-test was performed. All data represent mean ± SEM of 2-7 independent experiments. Related to **Figure S4**.

### Other cancer cell lines respond to CDK4/6 inhibitors and induce *F9* upregulation

To further determine if there is a wider implication for *F9* loss of function in other types of cancers we tested a panel of 22 cancer cell lines from different origins and molecular characteristic with increasing concentrations of Palbo, Abema and Ribo (**Figure 5A** **and Figure S5A**). As shown in **Figure 5B**, 8 cancer cell lines (including MCF7) responded to ≥ 2 CDK4/6 inhibitors in a dose-dependent and statistically significant manner (p < 0.05) (**Figure S5A**). Further validation of these 8 cells lines in a secondary screen re-evaluating the proliferation arrest induced by all three CDK4/6 inhibitors confirmed the response of 5 cell lines (p < 0.05) to more than two CDK4/6 inhibitors (MCF7, SKMEL28, ACHN, HT-29, SNU387) (**Figure 5C, Figure S5B, S5C**). Next, we determined which of these 5 cell lines (excluding MCF7 which we already validated) induced an upregulation of endogenous *F9* mRNA levels upon the treatment with different CDK4/6 inhibitors by qPCR. Of all the cancer cell lines analysed, the renal adenocarcinoma cell line (ACHN) was the only cell line to upregulate *F9* with Palbo and Abema (**Figure 5D**), while in the human colorectal adenocarcinoma cell line (HT29) and the hepatocellular carcinoma cell (SNU-387) induced endogenous levels of *F9* mRNA expression only with Palbo (**Figure 5D**). In accordance with our previous results, *F9* was not upregulated by Ribo in any of the cell lines analysed (**Figure 5D**). Interestingly, we show a partial proliferation bypass by crystal violet staining in ACHN upon shF9#4 expression (**Figure 5E**). This bypass could implicate that *F9* is only partially important in Palbo induced senescence in ACHN cells and that other mechanisms might be implicated. Altogether, our data highlight a partial relevance for *F9* mRNA expression in other cancer cell lines.

**Figure 5.**
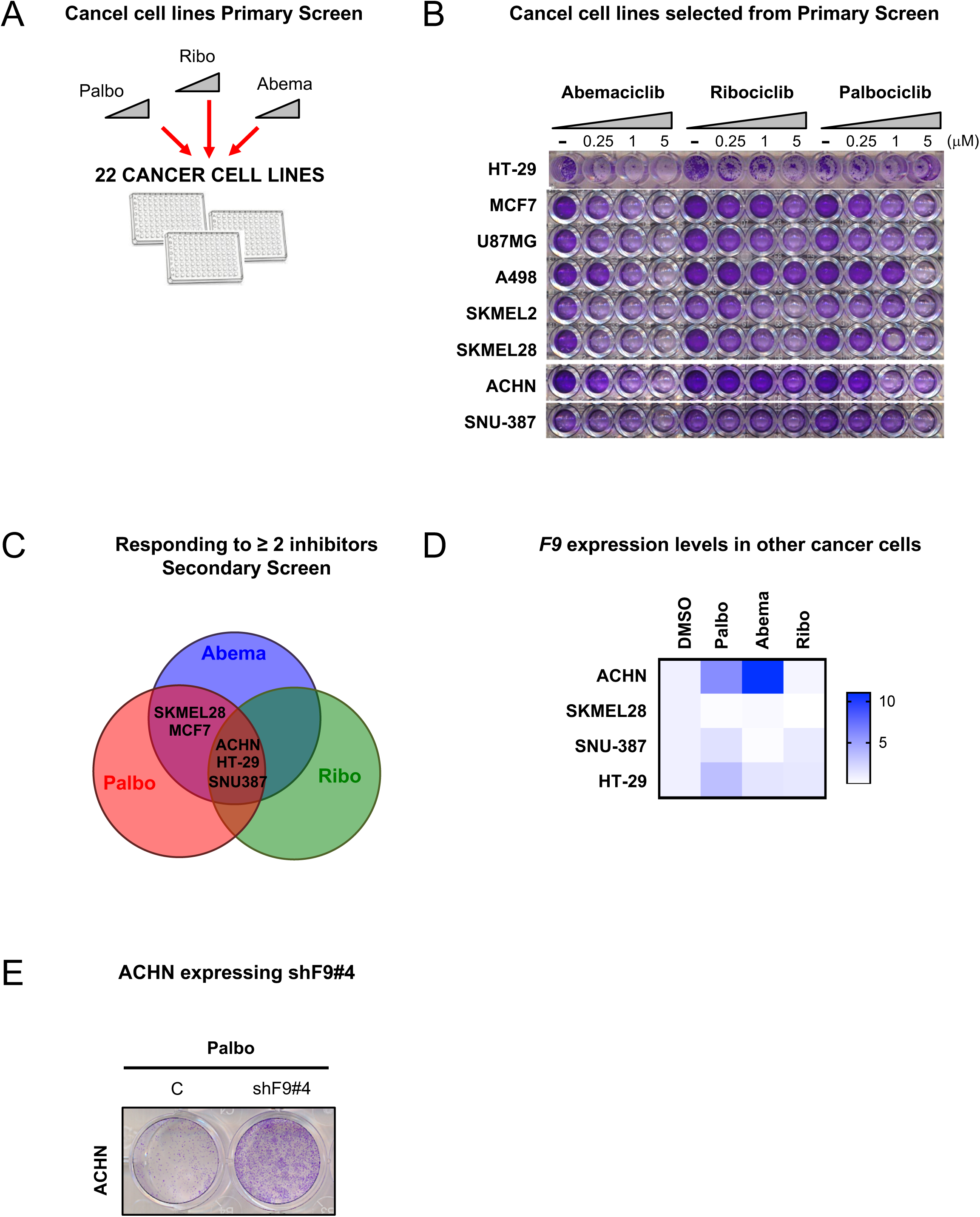
Response to CDK4/6 inhibitors in other cancer cell lines. **(A)** A panel of 22 cancer cell lines of different origins were treated with increasing concentrations of Palbo, Ribo and Abema. **(B**) Crystal violet staining showing the 8 cancer cell lines that responded in a statistically significant (p<0.05) and dose-dependent manner to more than 2 inhibitors. Representative experiment is shown. **(C)** Venn diagram shows that SKMEL28 (melanoma), MCF7 (breast cancer), ACHN (renal adenocarcinoma), HT-29 (colon) and SNU-387 (liver) cancer cell lines responded to two or more CDK4/6/ inhibitors (p<0.05) in a Secondary Screen and were selected for further validation. **(D)** Heat map showing *F9* mRNA expression in the indicated cancer cells after treatment with different CDK4/6 inhibitors. The map represents the mean of 3-5 independent replicates. **(E)** ACHN control or expressing shF9#4 stained with crystal violet after 20 days treatment with Palbo show a partial proliferation bypass. Representative picture of 3 independent experiments. See also **Figure S5**.

### *F9* is highly expressed in the tumour stroma

As we can observe an increase in *F9* mRNA levels in HFFF2 fibroblasts undergoing senescence, we next sought to explore published cancer datasets analysing tumour stroma^39^. We found that *F9* is upregulated in the tumour stroma in comparison with healthy stroma in breast and colon cancer, but not in prostate cancer lesions (**Figure 6A**). Interestingly, when analysing other transcripts implicated in the intrinsic coagulation pathway, where F9 is involved, most are also upregulated in breast cancer, while transcripts within the extrinsic pathway are mainly downregulated (**Figure 6B**)^39^. These data are in accordance with published studies showing a correlation between thrombosis and cancer^40,41^. As the tumour microenvironment is composed of cancer cells and stroma, we next sought to identify whether a cross talk existed between HFFF2 fibroblasts undergoing senescence by iRAS expression and MCF7 treated with DMSO or Palbo (**Figure 6C**). Senescence was induced in HFFF2 iRAS cells by treating them with or without 200nM 4OHT for 3 days, washed to prevent transfer of 4OHT, incubated with fresh media after which the conditioned media (CM) was collected (**Figure 6C**). Next, MCF7 pre-treated with or without 500nM Palbo were incubated with the CM from HFFF2 cells for 72h and we determined if we could observe changes in *F9* mRNA levels (**Figure 6D**). Interestingly, we found a sharp increase in *F9* mRNA levels when senescent MCF7 cells (treated with Palbo) were incubated with the CM of senescent HFFF2 (+4OHT). This increase also correlated with other SASP mRNA transcripts (**Figure 6E**) suggesting that senescent fibroblasts are reinforcing the SASP in already senescent MCF7 cells.

**Figure 6.**
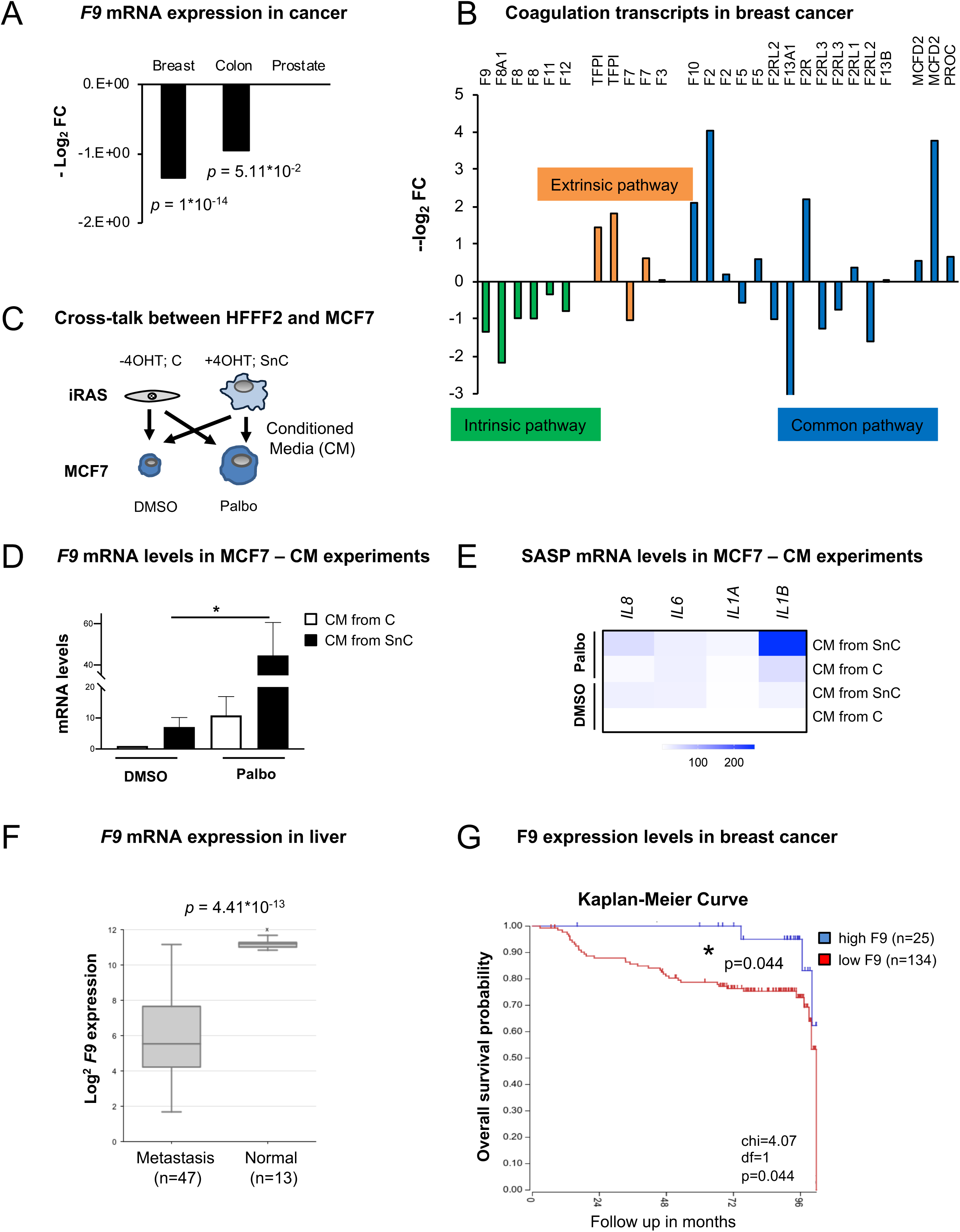
*F9* expression is important in different cancer types. **(A)** *F9* mRNA levels in tumour stroma *vs* healthy stroma in breast, colon and pancreatic cancers. Data are represented as -log_2_ fold change (FC) from^39^. Breast (n=12 samples from normal *versus* n=111 from tumour); colon (n=4 samples from normal *versus* n=13 from tumour); prostate (n=10 samples from normal versus n=8 from tumour). T-test Student analyses is performed. **(B)** mRNA expression levels of different mRNA transcripts from genes implicated in the intrinsic (green bars), extrinsic (orange bars) and transcripts common to both pathways (blue bars) in breast cancer. Comparison of tumour stroma (n=111) *vs* healthy stroma (n=12)^39^. **(C)** Schematic representation of panels **(D)** and **(E)**. MCF7 cells pre-treated with DMSO or 500nM Palbo were incubated for 72h with the conditioned media (CM) of control (−4OHT) (C) or senescent (+4OHT) (SnC) iRAS HFFF2 primary fibroblasts. Senescence was induced with 200nM 4OHT for 3, washed, incubated with fresh media and collected after 3 days. **(D)** *F9* mRNA levels and **(E)** heatmap for other SASP transcripts in MCF7 pre-treated with DMSO or 500nM Palbo and incubated with the CM from iRAS (-/+ 200nM 4OHT; C or SnC) for 72h. Data show the mean ± SEM of 4 independent replicates for *F9* and the mean of 3-6 independent experiments for the SASP. Two-way ANOVA with Dunnett’s multiple comparisons analyses was performed. **(F)** F9 expression levels in liver from normal (n=13 samples) and metastatic (n=47) liver. One- Way ANOVA analysis was performed to determine statistical significance. Dataset was calculated using R2. **(G)** Kaplan-Meier survival curve for high (blue) (n=25) or low (red) (n=134) F9 expression levels and overall survival prognostic in breast cancer. Chi-square = 4.07; p = 0.04 ^43^. Dataset calculated with R2.

### *F9* loss of function is associated with metastasis and worst survival prognostics in cancer

Our data suggest that increased levels of *F9* upon the induction of senescence maintain certain cancer cells in a stable cell cycle arrest, while *F9* loss of function promotes a proliferative advantage. In fact, in accordance with our data low levels of *F9* are associated with liver metastasis in comparison with normal liver^42^ (**Figure 6F**). Importantly, high levels of F9 expression are a sign of good prognostic for survival in breast cancer in comparison with those patients presenting low levels of F9 expression when analysing the breast cancer dataset^43^ (**Figure 6G**). Altogether, these data show the potential of using F9 levels as a biomarker for patient stratification not only to predict response to CDK4/6 inhibitors but also as a prognostic marker to determine overall survival in breast cancer.

## DISCUSSION

A better understanding of the mechanisms regulating senescence induced by treating cancer cells with CDK4/6 inhibitors is needed in order to increase the efficacy of targeted therapy in cancer and to be able to stratify patients. While previous *in vitro* studies showed that RB, cyclin D1 and p16 could predict Palbociclib response^44,45,46^, results from Phase II/III clinical trials show no correlation between the expression of CCND1, p16 or Ki67 leaving no prognostic or predictive biomarkers that allow to secure drug efficacy and diminish drug resistance or response^16,47,48^. In this study, we provide evidence that the coagulation factor IX (*F9*) plays an important role in regulating the cell cycle arrest and senescence phenotype induced by CDK4/6 inhibitors in ER^+^ breast cancer cell lines and other cancer cells.

CDK4/6 inhibitors (Palbociclib, Abemaciclib, Ribociclib) are considered highly selective new generation small molecule inhibitors that bind to the CDK4 and CDK6 ATP-binding pocket, leading to the inactivation of CDK4/6-CyclinD complexes with the subsequent inhibition of RB phosphorylation and induction of a G1 phase arrest^2^. In fact, CDK4/6 inhibitors induce a senescence-like state in different cell types^8,49,50^. However, their highest effect was demonstrated in preventing hormone-dependent cell-cycle entry in advanced ER^+^ breast cancer cells^7^. Our results confirmed that the treatment of the ER^+^ breast cancer cell line MCF7 with different concentrations of CDK4/6 inhibitors, induces a stable proliferation arrest and a senescence-like phenotype shown by an increase in β- galactosidase activity and the expression of senescence markers such as p21^CIP1^. While the three CDK4/6 inhibitors have demonstrated greater efficacy in combination with endocrine therapy in postmenopausal women (PALOMA-2, MONALEESA-2, MONARCH- 3 trials) and significantly prolonged the progression-free survival from 18 months to more than 27^16,51,52,53^, genes regulating senescence induction in cancer cells have not been identified yet. To gain further insight into the mechanisms inducing senescence by CDK4/6 inhibitors, we performed a genome wide CRISPR/Cas9 screen. This would allow us to identify genes whose loss of function prevented the cell cycle arrest induced by Palbo treatment in MCF7 cells, thus identifying genes that can predicted a lack of response to Palbo. We found a total of eighteen sgRNA enriched after two weeks of treatment, targeting genes which regulate PI3K or p53 signalling pathways, previously known to regulate senescence or senescence-related phenotypes^54,55,56,57^. However, KEGG analysis revealed an enrichment for genes whose loss of function regulated the blood coagulation pathway, suggesting that genes participating in the coagulation cascade (*F9*, *PROZ*) could regulate senescence induced by Palbociclib.

The identification of genes participating in the blood coagulation pathway, is in accordance with previous publications were it was demonstrated that inflammaging, hypercoagulability and cellular senescence share common pathways^58^. *F9* participates in the intrinsic pathway of blood coagulation by converting factor X in its active form regulating the haemostasis program^59^. Our results, demonstrate not only that *F9* levels are upregulated and secreted when MCF7 cells are treated with CDK4/6 inhibitors (specially Palbo and Abema) but also, when human primary fibroblasts are triggered to induce senescence by expressing the H-RAS^G12V^ or when treated with etoposide or Palbociclib. Furthermore, endothelial cells also express higher levels of *F9* upon senescence induction when using Palbociclib. Even though the link between senescence and haemostasis is not fully clear, *Wiley et al.* demonstrated that human fibroblasts undergoing senescence secrete a subset of haemostasis-related factors as part of the SASP^33^. It is however interesting to note that they did not find factors implicated in the coagulation pathways suggesting that maybe different triggers or cell cultures could favour either general haemostasis or the coagulation pathway specifically.

In concordance with our data, the combination of MEK and CDK4/6 inhibitors in pancreatic adenocarcinoma suppresses cell proliferation and induces the release of SASP factors enriched in pro-angiogenic proteins promoting tumour vascularization which, in turn, enhances drug delivery and efficacy^50^. In line with these findings, senescent human primary fibroblasts also release pro-angiogenic factors such as VEGF^60^. Altogether, these findings prompts us to think that *F9* and the coagulation pathway play an important role in senescence. In fact, the coagulation pathway has been shown to be upregulated in senescence^61^ and ageing^62,63^ by others. Furthermore, F9 has been associated with frailty ^64^ and is a genetic risk factor estimated to contribute to thrombosis incidence in the elderly^62,65^.

Although pro-senescence therapies are considered a potential anti-cancer strategy, the accumulation of senescent cells in the tissue is deleterious. In fact, the elimination of senescent cells promotes tumour clearance, tissue regeneration and ameliorates age- related pathologies^26,35,66^. This process is mainly controlled by the recruitment of the immune system through SASP factors^9, 11,28,67^ that, in the same line, are able to modify platelet function, resulting in increased efficacy^68^. Our data show that Palbociclib induces certain SASP factors in MCF7, but this secretion was partially prevented by *F9* loss of function. Although more experiments would be needed it is tempting to speculate a role for F9 as part of the SASP in recruiting immune cells or regulating platelet function in this context. In fact, CDK4/6 inhibitors have been shown to enhance T-cell activation^69^ and NK recruitment^67^ although a direct role for F9 has not been described. Interestingly, it has been show that hepatocyte-derived human coagulation F9 expression can induce regulatory CD4^+^ T cells in mice^70^ suggesting a role for F9 in regulating immunosurveillance.

It is well established that a link between thrombosis and cancer exists^41,71^. The expression of the oncogene *MET* in the liver *in vivo* not only causes hepatocarcinogenesis but also blood hypercoagulation and fatal internal haemorrhage bleeding^40^. In fact, the use of antiplatelet therapy in liver cancer prevents cross-talk between platelets and immune cell interaction^72^. Curiously, chemotherapy treatment increases the risk of thrombosis^73^ and a link showing that senescence promotes the adverse effects of chemotherapy and cancer relapse has been recently shown^29^. It would therefore be interesting to further explore the implication of the coagulation pathway and F9 in particular in the context of chemotherapy and senescence.

In summary, here we provide evidence for the involvement of two different genes within the coagulation pathway, *F9* and *PROZ*, in regulating senescence in different contexts. Our results show that *F9* is partially responsible for CDK4/6 inhibitors response in breast and renal carcinoma *in vitro*. Importantly, we unveiled a correlation between the levels of F9 and cancer progression studying different cancer related datasets. Altogether, we believe *F9* could be considered as a potential biomarker to predict CDK4/6 inhibitors response when used as first line treatment in cancer.

## EXPERIMENTAL PROCEDURES

### Cell culture

All cancer cell lines used in this study were obtained from American Type Culture Collection (ATCC). Human foreskin fibroblasts (HFFF2) were obtained from Culture Collections (Public Health England, UK). All cell lines were grown in high glucose Dulbecco’s Modified Eagle Medium (DMEM) (Gibco) except for HCT-116, SK-OV-3, Capan2 and HT29 that were grown in McCoys 5a medium, SNU-387, NCI-H23, OVCAR- 3 that were grown in RPMI and A549, PC-3 that were grown in F12K medium. All media was supplemented with 10% foetal bovine serum (FBS) (Thermo Fisher) and 1% of antibiotic-anti-mycotic (Thermo Fisher). Human umbilical vein endothelial cells (HUVEC) from pooled donors were purchased from Promocell (Heidelberg, Germany). The cells were grown in M199 (Life Technologies, Grand Island, NY) with 20% fetal bovine serum (Labtech, Heathfield, UK), 10U/ml heparin (Sigma, St. Louis, MO), and 30μg/ml endothelial cell growth supplement (Sigma). Experiments were performed using HUVECs between passage 3 and 5.

### Senescence induction

MCF7 cells were treated with different concentrations (100nM, 200nM, 500nM or 1000 nM) of Palbociclib (PD0332991) (APExBIO), Abemaciclib (LY2835219) (Selleckchem) or Ribociclib (LEE011) (Selleckchem) and the other cancer cell lines with the indicated concentration of inhibitor. Human primary fibroblasts (HFFF2) were treated with 50μM of Etoposide (Sigma-Aldrich) for 2 days and cells collected at day 7 or treated with 1μM Palbociclib for 7 days. HFFF2 expressing pLNC-ER:RAS vector, were induced to senescence by adding 200nM 4-hydroxytamoxifen (4OHT) (Sigma-Aldrich) for 6 days. All treatments were done using DMEM supplemented with 10% FBS and 1% of antibiotic-anti- mycotic. HUVEC cells were treated with 500nM Palbo for 7 days to induce senescence.

### F9 recombinant experiments

Recombinant experiments were carried out by using F9 recombinant protein (R&D systems). MCF7 cell line was treated with 10 μg/ml in complete medium for two rounds of 72 hours (total of 6 days treatment). At the end of the experiment cells were fixed with 4% paraformaldehyde (PFA) and used for immunofluorescence studies.

### Retroviral and lentiviral infections

The generation of stable retroviral and lentiviral expression was carried out following previous studies^8, 34,35,74^. Briefly, retroviral particles were generated by transfecting pLNC-ER:H-RAS^G12V^ plasmid and retroviral helper plasmids (vsvg and gag-pol) with Polyethylenimine (PEI) in HEK293T packaging cells for 48h. Recombinant lentiviral particles were generated using the second-generation packaging vectors psPAX2 and pMD2.G using PEI in HEK293T. The supernatant containing retrovirus or lentivirus was then filtered with 0.45µm filters (Starlab) and applied to HFFF2 cells in the presence of 4µg/ml polybrene (hexadimethrine bromide; Sigma-Aldrich) following 3 rounds of infection. Cells were subsequently selected with the appropriate antibiotic resistance either 0.5μg/ml puromycin or 300μg/ml neomycin (Invitrogen). For lentiviral infections with the sgRNA a pool for the 4 sgRNA (5μg DNA per single sgRNA) targeting a single gene was generated by transfecting equal amounts of DNA and the packaging vectors psPAX2 and pMD2.G. Infection was performed as described earlier for lentivirus.

### Genome-wide CRISPR/Cas9 library amplification and sequencing

The Human CRISPR knockout Pooled Library (GeCKO*v2*) was purchased from Addgene (#1000000048) and amplified using *E. coli* competent cells. The library contains 123,411 unique sgRNA sequences targeting 19,050 genes within the human genome. After amplifying the library as described^31,75^, viral particles were produced in HEK293T cells and MCF7 cells were infected at a multiplicity of infection (MOI) of 0.2 - 0.5 following the lentiviral protocol previously described^31,75^. Cells were selected with puromycin (1μg/ml) for 72h after the GeCKO library infection. After selection, cells were plated at low density and treated with 200nM Palbociclib for 14 days to either determine proliferation or sent DNA for genomic DNA sequencing. Crystal violet staining was used to assess cell proliferation and determine the library bypass efficacy. After MCF7 cell infection and selection with the GeCKO library, genomic DNA was extracted at days 0 and day 14 after infection using the QIAmp Blood and Cell Culture DNA midi kit (Qiagen). The PCR was performed by QMUL Genome Centre.

The accession number for the sequencing data reported in this paper is GEO: XXXXXXXX.

### CRISPR sgRNA generation

The online guide design tool (http://crispr.mit.edu) was used to identify sgRNA sequences. The highest scoring guides were selected. Primers for the sgRNA sequences were ordered and the complementary sequences annealed at 37C for 30min, followed by incubating the annealed primers at 95C for 5 min and then ramped down to 25C at 5C degrees per min. The annealed synthetic sgRNA oligonucleotides were cloned into pLentiCRISPR*v2* vector (Addgene #52961) at BsmBI restriction sites. The sgRNA sequences used in this study are:

sgF9#1 GCAGCGCGTGAACATGATCATGG

sgF9#2 CACTGAGTAGATATCCTAAAAGG

sgF9#3 ATGATCATGGCAGAATCACCAGG

sgF9#4 CTAAAAGGCAGATGGTGATGAGG

sgLPAR5#1 CCCAGAGGGCTAGCGCGTTGAGG

sgLPAR5#2 CCAGAGGGCTAGCGCGTTGAGGG

sgLPAR5#3 GGAAGATGGCGCCCGTCGTCTGG

sgLPAR5#4 GCGTAGTAGGAGAGACGAACGGG

sgPROZ#1 TGAGGGCTCCACACGATGGAGGG

sgPROZ#2 GGTCCTCGCCCTCCATCGTGTGG

sgPROZ#3 CTGAGGGCTCCACACGATGGAGG

sgPROZ#4 GCTCCACACGATGGAGGGCGAGG

sgMOGAT#1 CCGCAATGTAGTTCCGAGAGGGG

sgMOGAT#2 GCCCGCAATGTAGTTCCGAGAGG

sgMOGAT#3 GTTCCGCAGTAACAGCGTGAAGG

sgMOGAT#4 GCTGTTACTGCGGAACCGAAAGG

sgRB#3 GGTGGCGGCCGTTTTTCGGGGGG

sgRB#4 CGGCGGTGGCGGCCGTTTTTCGG

To identify positive clones the primer hU6 CRISPR 5’-GAGGGCCTATTTCCCATGATT- 3’ was used in combination with the reverse primer for each specific clone and isolated clones were SANGER sequenced.

### Cell proliferation experiments

For cell proliferation studies, 100 cells were plated in each well of a 96-well plate and treated with CDK inhibitors (Palbociclib, Abemaciclib, Ribociclib) for 0, 6, 15, 20 days. The medium was replaced every other day either with drug or drug-free medium. For replating experiments, cells were treated with Palbociclib for 6 days, counted and replated at low density in a 96 well plate until day 20. Drug-withdrawal was performed by treating the cells 6 days in the presence of Palbociclib and removing the drug from day 6 to day 20. Cells were then fixed, stained with 0.5% crystal violet, solubilized with 30% acetic acid solution and absorbance was measured at 570nm.

For colony formation assay, 5000 cells plated in each well of a six-well plate, treated with CDK inhibitors 20-30 days (once control reaches confluence). Cells were then washed with PBS, stained with crystal violet and scanned to obtain the pictures.

### CDK4/6 inhibitors dose-response studies

Dose-response studies were carried out using the panel of cancer cell lines listed in the Cell Culture section. Cells were plated in a 96 well plate at 500-1000 cells/well (based on seeding density calculations) and treated with increasing concentrations (0.25 - 5 μM) of Palbociclib, Abemaciclib and Ribociclib. Medium containing the drugs or DMSO was replaced every other day during 10 days. The plates were then stained with 0.5% crystal violet solution and scanned. Crystal violet quantification was performed by solubilizing crystal violet staining with 30% acetic acid and measuring the absorbance at 570 nm.

### β-galactosidase staining

Cells were washed with PBS and fixed with 0.05% (w/v) glutaraldehyde (in PBS) for 15 min at room temperature. Cells were washed a second time with PBS and incubated with 5-bromo-4-chloro-3-indolyl-beta-D-galacto-pyranoside (X-gal) solution for 1h at 37°C. Cells were imaged after 12 - 24h using a light microscope (Nikon) at 20X magnification and single representative images of each well were taken. Fluorescent β-Galactosidase was performed according to the manufacturer’s instructions using the following commercial kit (Sigma-Aldrich, #F2756). Briefly, 33μM of the β-gal substrate C_12_FDG (Fluorescein di-β-D-galactopyranose) (F2756 Sigma-Aldrich) was added to the cells for 8h at 37°C, After, the cells were washed with PBS and fixed with 4% PFA.

### RNA extraction, cDNA synthesis and qPCR

Total RNA was extracted using TRIzol Reagent (ThermoFisher) according to the manufacturer’s instructions. cDNA synthesis was performed using High Capacity cDNA Reverse Transcriptase kit (ThermoFisher). qPCR reactions were performed using SYBR Green PCR Master Mix (Applied Biosystems) on a 7500 Fast System RealTime PCR cycler (Applied Biosystems). Primer sequences used in this study are:

F9 Forward 5’-CAGTGTTCAGAGCCAAGCAA-3’

F9 Reverse 5’-CATGGTGAACACGAAACCAG-3’

PROZ Forward 5’-CACCCCTGAGAAAGACTTCG-3’

PROZ Reverse 5’-GGAGCCTCTGTGTTCTCTGG-3’

RB Forward 5’-AACCCAGGAAGGAATGGCT-3’

RB Reverse 5’-CTGCGTTCAGGTGATTGATG-3’

IL8 Forward 5’-GAGTGGACCACACTGCGCCA-3’

IL8 Reverse 5’-TCCACAACCCTCTGCACCCAGT-3’

IL6 Forward 5’-CCAGGAGCCCAGCTATGAAC-3’

IL6 Reverse 5’-CCCAGGGAGAAGGCAACTG-3’

CCL20 Forward 5’-GGCGAATCAGAAGCAGCAAGCAAC-3’

CCL20 Reverse 5’-ATTGGCCAGCTGCCGTGTGAA-3’

IL1A Forward 5’-AGTGCTGCTGAAGGAGATGCCTGA-3’

IL1A Reverse 5’-CCCCTGCCAAGCACACCCAGTA-3’

IL1B Forward 5’-TGCACGCTCCGGGACTCACA-3’

IL1B Reverse 5’-CATGGAGAACACCACTTGTTGCTCC-3’

CDKN1A Forward 5’-CCTGTCACTGTCTTGTACCCT-3’

CDKN1A Reverse 5’-GCGTTTGGAGTGGTAGAAATCT-3’

ACTIN Forward 5’-GCCCTGAGGCACTCTTCCA-3’

ACTIN Reverse 5’-CGGATGTCCACGTCACACTTC-3’

RSP14 Forward 5’-CTGCGAGTGCTGTCAGAGG-3’

RSP14 Reverse 5’-TCACCGCCCTACACATCAAACT-3’

### Protein lysis and western blot

Cells were lysed in ice-cold Lysis Buffer 6 (R&D systems) supplemented with 10 μL/mL of protease inhibitor cocktail. Total protein content was determined by Precision Red Reagent (Sigma) protein assay. Twenty micrograms of total protein were separated on 10% SDS- PAGE and transferred to a polyvinylidene fluoride (PVDF) membrane (Millipore Co., Bedford, MA). Protein transfer was checked by staining the membrane with Ponceau S red (Sigma-Aldrich). The membrane was then blocked using 5% bovine serum albumin (BSA) (Sigma) or 5% milk (Sigma) in PBS supplemented with 0.05% Tween-20 (Sigma) (PBST). Primary antibodies RB1 (BD; Cat# 554136) and β-Actin (Abcam, Cat# ab8226) were incubated overnight at 4C. After four washes with PBST, the membrane was incubated with a secondary antibody for 1h at room temperature. Protein bands were detected using SuperSignal West Pico PLUS Chemiluminescent Substrate (Thermo Fisher Scientific) using the ChemiDoc XRS+ System (Bio-Rad).

### Immunofluorescence

Cells were grown in a 96-well plate and fixed with 4% paraformaldehyde for 10 min at RT. Cells were then washed twice with PBS and permeabilized by incubating with 0.4% Triton X-100 (xx) in PBS for 10 min at RT. After a PBS wash, cells were blocked 30 min at RT using 1% BSA in PBS supplemented with 0.1% Tween-20 (PBST). Primary antibodies (details found at the end of the section) were diluted in 1% BSA-PBST and incubated overnight. For BrdU staining, cells were treated with DNaseI and MgCl_2_ simultaneously with the primary antibody. Cells were then washed with PBS and incubated with their respective secondary antibody for 1h at RT. Nuclei were stained with DAPI (Sigma-Aldrich). Images were acquired using INCell 2200 automated 991 microscope (GE) and INCell 2200 Developer software version 1.8 (GE) was used for image analysis. Antibody used in this study are: p21^CIP^ (Abcam, Cat# ab109520), Ki67 (Abcam, Cat# ab92742), BrdU (Abcam, ab6326; 1:500) and F9 (Proteintech, Cat# 21481-1-AP).

### Conditioned media experiments

Donor cells (HFFF2 iRAS) were treated in the presence or absence of 200nM 4- hydroxytamoxifen (4OHT) (Sigma-Aldrich) for 3 days, washed and replenished with fresh media to prevent carrying the 4OHT. Conditioned medium (CM) was collected and supplemented to 10% FBS and added to MCF7 recipient cells for 72h. MCF7 cells were pre-treated in the presence or absence of 500nM Palbociclib prior to adding the CM.

### Statistics

Dataset analysis were performed using R2: Genomics Analysis and Visualization Platform (http://r2.amc.nl). STRING interactions were identified using the functional protein association networks (https://string-db.org/). Kyoto Encyclopedia of Genes and Genomes (KEGG) pathway analysis was performed using Panther pathway analysis (http://www.pantherdb.org). A p < 0.05 is considered significant throughout the paper as follows: *p<0.05; **p<0.01; ***p<0.001.

## AUTHOR CONTRIBUTIONS

A.O. conceived and designed the study. P.C.F. performed most of the experiments with the help of O.E. and J.F.L. M.B. amplified the sgRNA library and carried out the GeCKOv2 whole genome wide screen. T.P.M. and T.D.N. performed the endothelial cells experiments. M.D.M. provided reagents. P.C.F. and A.O. designed the experiments and wrote the paper. All the authors discussed the results and commented on the manuscript.

## ACKNOWLEDGEMENTS

We are extremely grateful to Belen Pan-Castillo for very useful scientific discussions on the project. We want to thank the Queen Mary University of London Genome Centre, Dr. Luke Gammon and Dr. Gary Warnes for technical support. This paper was funded by the BBSRC (BB/P000223/1), The Royal Society (RG170399) and Barts Charity (MGU0497) grants to A.O. M.M. was funded by PI19/00145 from the Health Institute ‘Carlos III’ (ISCIII, Spain). P.C.F. and J.F.L. were funded by Xunta de Galicia (INB606B 2017/014 and ED481B 2017/117 respectively). M.B. was funded by MRC (MR/K501372/1). T.P.M. was funded by a QMUL funded PhD programme and T.D.N.’s lab is currently funded by a Barts Charity project grant (MGU0534). P.C.F. is currently funded by GAIN (IN607B2020/12) Xunta de Galicia.

## CONFLICT OF INTEREST

A.O. forms part of the Starklabs Scientific Advisory Board and has an unrelated project funded by Starklabs.

---

**Figure S1.**
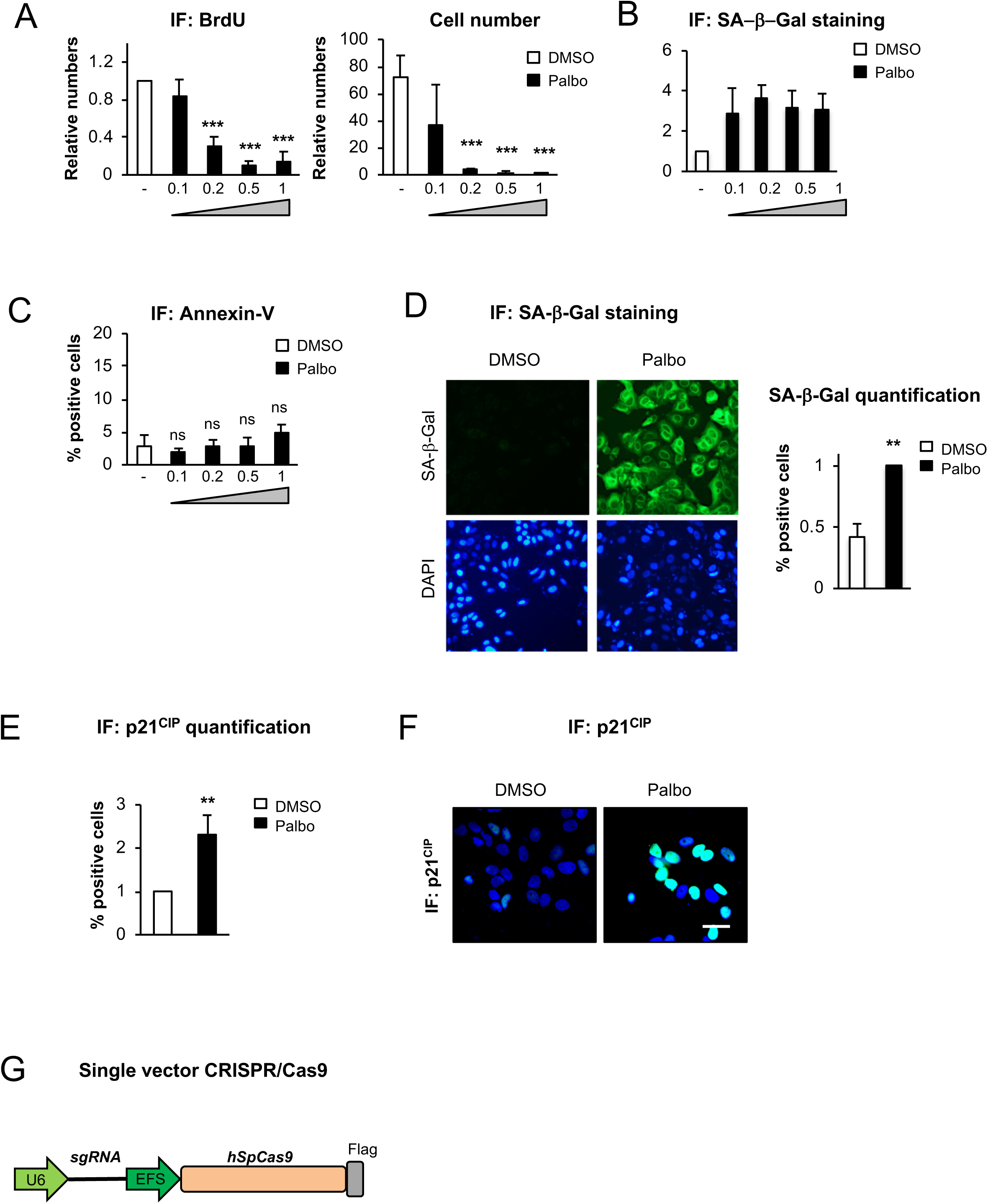
Palbociclib induces a senescent-like phenotype in MCF7 breast cancer cell line. **(A)** Quantification of BrdU incorporation and relative cell number in MFC7 treated with different concentrations (0.1, 0.2, 0.5 and 1 μM) of Palbociclib (Palbo) for 14 days. **(B)** SA-β-galactosidase (SA-β-Gal) staining in MCF7 cells treated with different concentrations of Palbo for 7 days. **(C)** The graph represents the percentage of Annexin V positive cells after 14 days Palbo treatment with different concentrations. **(D)** Representative images (left panel) and quantification (right panel) for SA-β-Gal staining in MCF7 cells treated with 200nM Palbo for 14 days. Graph shows the mean ± SEM of 4 independent experiments. Two-tailed Student’s t-test was used to calculate statistical significance. **(E)** Quantification and **(F)** representative pictures of the percentage of p21^CIP^ positive cells upon 14 days 200nM Palbo treatment. Two-tailed Student’s t-test was used to calculate statistical significance. Scale bar: 50 μm. **(G)** Diagram for the lentiCRISPR*v2* one vector system used in the GeCKO library and to clone individual sgRNAs. The plasmid contains an expression cassette for human Cas9 (*hSpCas9*) and the sgRNA in the same vector. Related to Figure 1.

**Figure S2.**
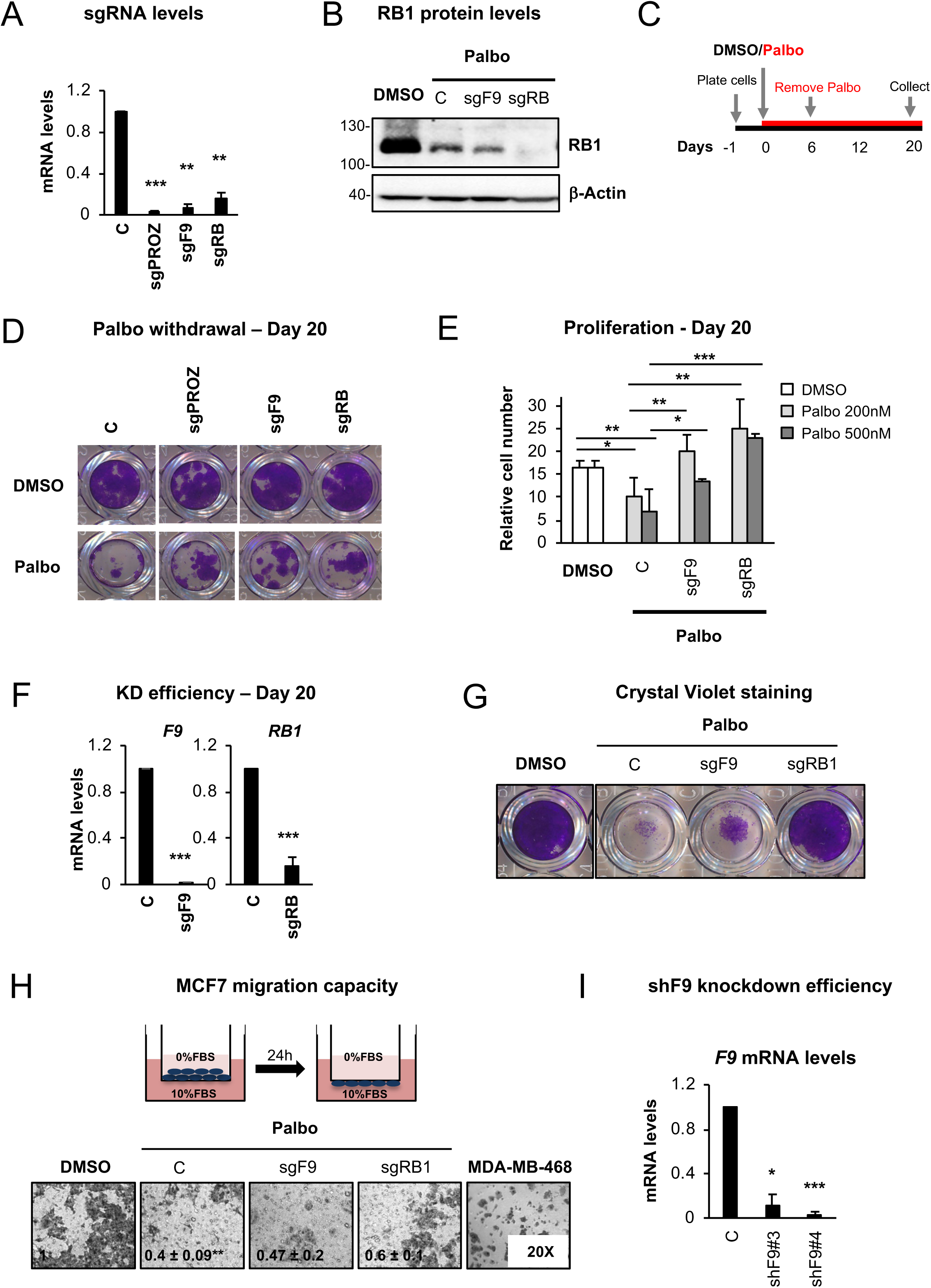
Validation of sgRNA efficiency and proliferative advantage in MCF7 cells. **(A)** Relative mRNA expression levels of *PROZ*, *F9* and *RB* in MCF7 after their respective sgRNA infection and selection. Graph shows the mean ± SEM of 2 independent experiments. Two-tailed Student’s t-test was used to calculate statistical significance compared to the Control sample. **(B)** Representative western blot showing RB knockout upon sgRB expression in MCF7 cells. β-actin was used as a loading control. Representative blot for 4 independent experiments. **(C)** Timeline and strategy followed to confirm that the senescence proliferative arrest is maintained after Palbo is removed and washed out. MCF7 cells were treated with DMSO or 500nM Palbo for 6 days, after which the drug was removed and the cells were grown and collected at day 20. **(D)** Crystal violet staining showing the proliferation rate of MCF7 cells expressing sgRNAs (sgF9 and sgRB) at day 20. Palbo was removed after day 6 to confirm the induction of a stable cell cycle arrest. A representative experiment of 3 independent experiments is shown. **(E)** Relative cell number quantification in MCF7 control or expressing sgRNAs (sgF9, sgRB) using 200nM (light grey) or 500nM (dark grey) Palbo treatment for 20 days. Data represent the mean ± SEM of 3 independent experiments. **(F)** mRNA levels for *F9* and *RB1* by qPCR in MCF7 cells 20 days after setting the experiment. Data represent the mean ± SD of 3 independent experiments. Two-tailed Student’s t-test was used to calculate statistical significance. **(G)** Crystal violet staining for MCF7 expressing sgF9 and treated with 500nM Palbo. A representative experiment is shown. sgRB is used as positive control. **(H)** 24h MCF7 migration assays upon 500nM Palbo treatment for 20 days expressing sgF9 and sgRB. MDA-MB-468 cells were used as a positive control. Representative pictures of 3 independent experiments is shown. Data show the mean ± SEM of 3 independent experiments. One-Way ANOVA with Dunnett’s multiple comparisons to Palbo was performed. **(I)** qPCR to confirm the efficacy of two independent shRNA targeting F9 (shF9#3 and shF9#4). Data represent the mean ± SEM. Two tailed students t-test analysis was performed. Related to Figure 2.

**Figure S3.**
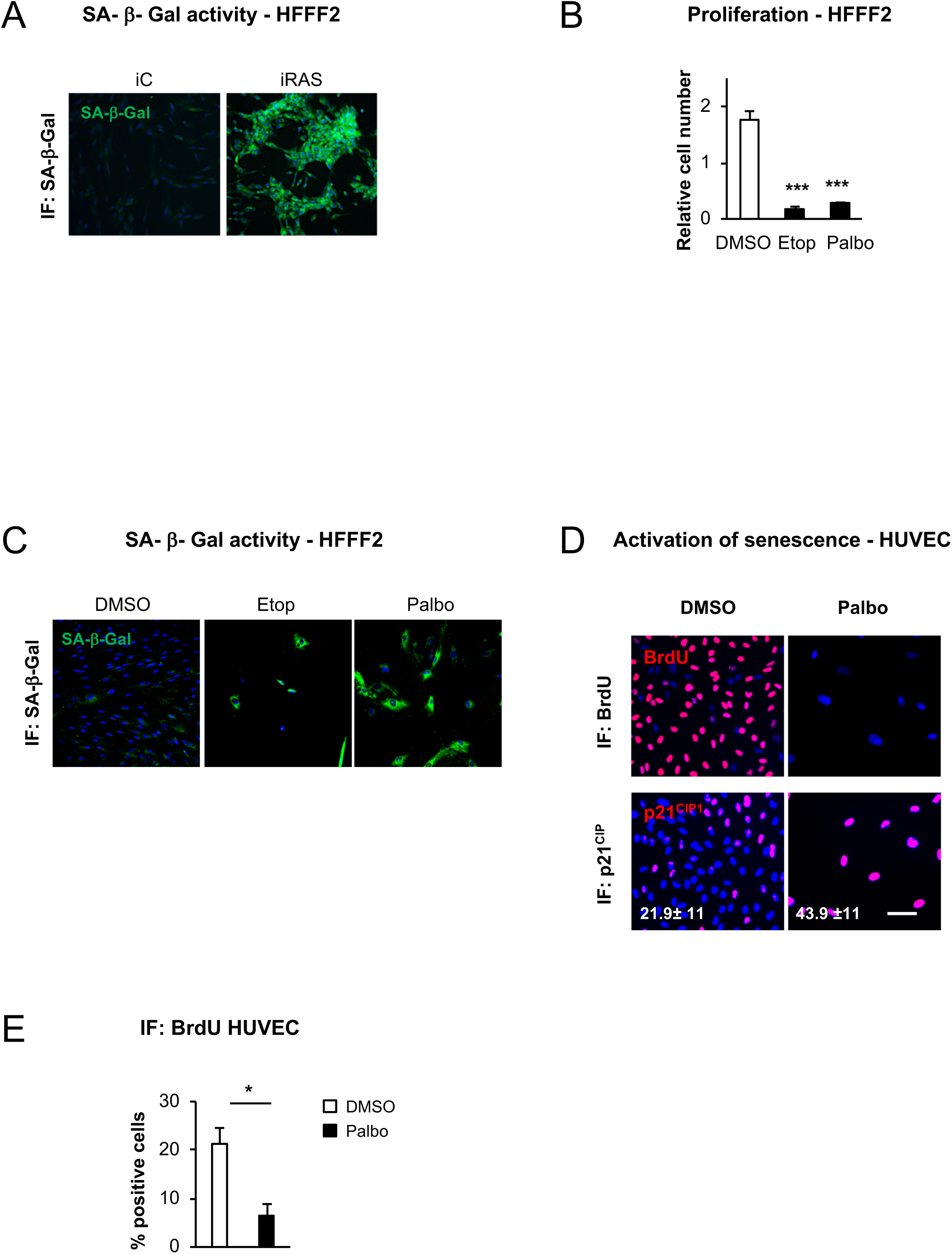
Induction of senescence in human primary fibroblasts (HFFF2) and primary endothelial cell (HUVEC) cultures. **(A)** Representative images of SA-β- galactosidase activity (SA-β-Gal). Human primary fibroblasts (HFFF2) expressing an empty vector ER:EV (iC) or ER:H-RAS^G12V^ (iRAS) were treated with 200nM 4OHT for 6 days to induce senescence. SA-β-Gal is shown by the incorporation of the fluorescent compound C_12_FDG (green). **(B)** HFFF2 relative cell number after 2 days treatment with 50 μM of Etoposide followed by 5 days with fresh media or 7 days with 1μM Palbociclib. Data show the mean ± SEM of 3 independent experiments. Two-tailed students t-test was performed compared to Control sample. **(C)** HFFF2 treated with 50μM of Etoposide for 2 days followed by 5 days with fresh media and 1μM Palbo for 7 days were incubated for 8h with C_12_FDG compound. SA-β-gal activity (green) was determined by fluorescent signal and representative images are shown. **(D)** Representative IF images for p21^CIP1^ and BrdU in HUVEC (human umbilical vein endothelial cells) control or treated with 500nM Palbo for 7 days. Representative images are shown from 3-4 independent experiments. Scale bar: 50 μm. **(E)** The graph represents the quantification for the percentage of HUVEC cells staining positive for BrdU. The data represent the mean ± SEM of 3 independent experiments. Two-tailed Student’s t-test was used as test. Related to Figure 3.

**Figure S4.**
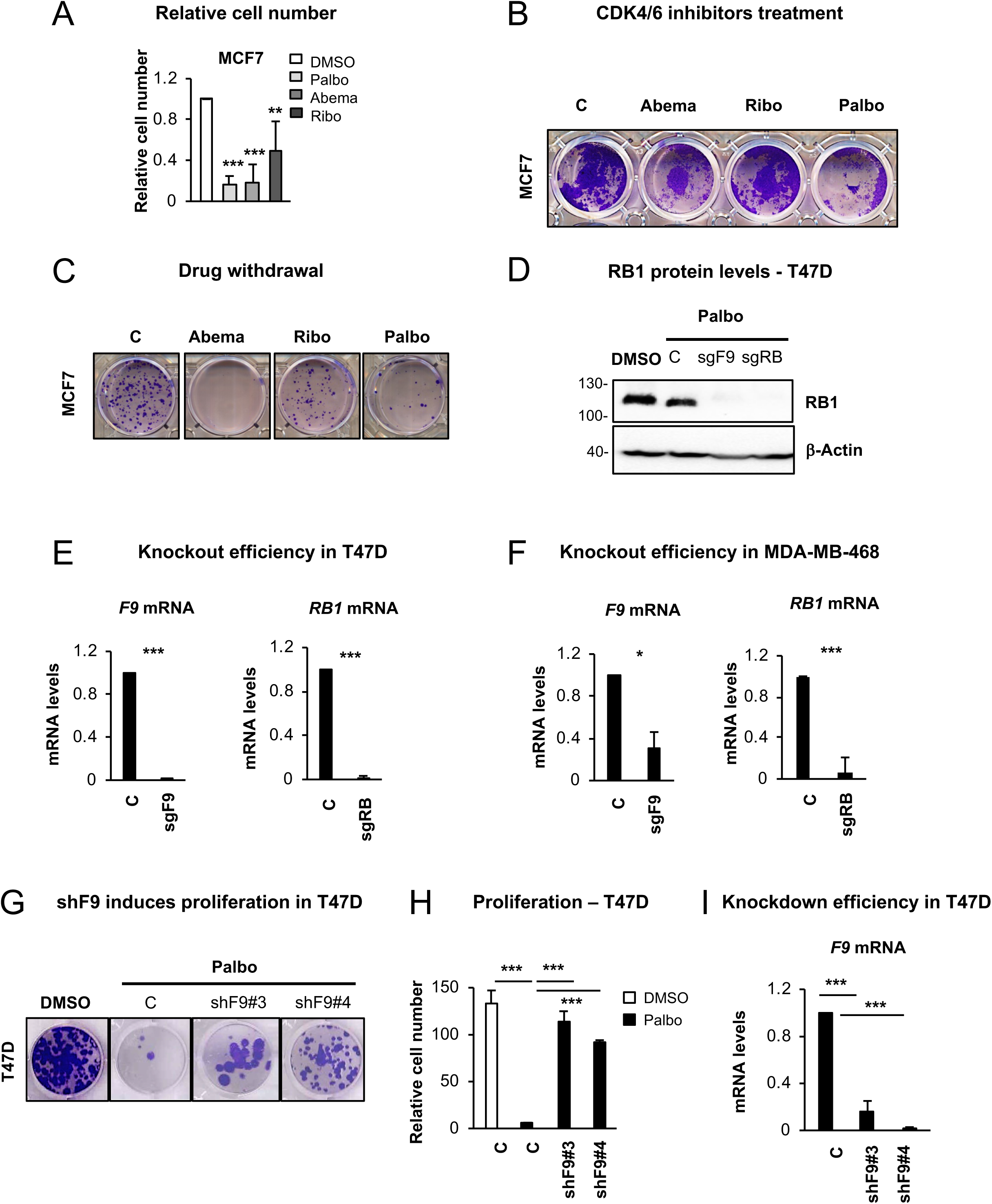
CDK4/6 inhibitors response in a variety of cancer cell lines. **(A)** Relative cell number quantified after MCF7 cells were treated with 1μM of different CDK4/6 inhibitors (Palbociclib, Abemaciclib, Ribociclib). Data show the mean ± SD of 2-5 independent experiments. Two-tailed student’s t-test analysis was performed. **(B)** Clonogenic assay shows the proliferation rate of MCF7 cells after 14 days treatment with 1μM Abema, 1μM Palbo and 1μM Ribo. Representative experiment of 2 biological replicates. **(C)** Crystal violet staining showing the effect on proliferation for after 6 days treatment with Palbo, Abema and Ribo. Experiment was stopped 20 days after treatment. Representative experiment is shown. **(D)** Representative western blot showing RB knockout in T47D cells. β-actin was used as a loading control. Blot representative of 3 independent experiments. **(E)** *F9* and *RB1* mRNA levels were determined by qPCR in T47D cells expressing sgF9 or sgRB1. Data show the mean ± SEM of 3 independent experiments. Two-tailed student’ t-test analysis was performed. **(F)** Knockout efficiency for sgF9 and sgRB in MDA-MD-468 cells. mRNA levels were determined by qPCR. Data show the mean ± SEM of 3 biological replicates. Two-tailed student’s t-test analysis was performed. **(G)** Crystal violet staining showing the proliferative rate of T47D cells expressing two independent shRNA targeting F9 (shF9#3 and shF9#4) treated with 1μM Palbo for 20 days. Representative experiment of 3 biological replicates. **(H)** Proliferation rate of T47D expressing shF9#3 and shF9#4 treated with 1μM Palbo for 20 days. The data represent the mean ± SEM of 7 independent experiments. One Way ANOVA with Dunnett’s multiple comparisons to Palbo C sample was performed. **(I)** qPCR analysis for the levels of *F9* mRNA in T47D cells expressing shF9#3 and shF9#4. Data show the mean ± SEM of 3 independent experiments. Two-tailed student’ t-test analysis was performed. See also to Figure 4.

**Figure S5.**
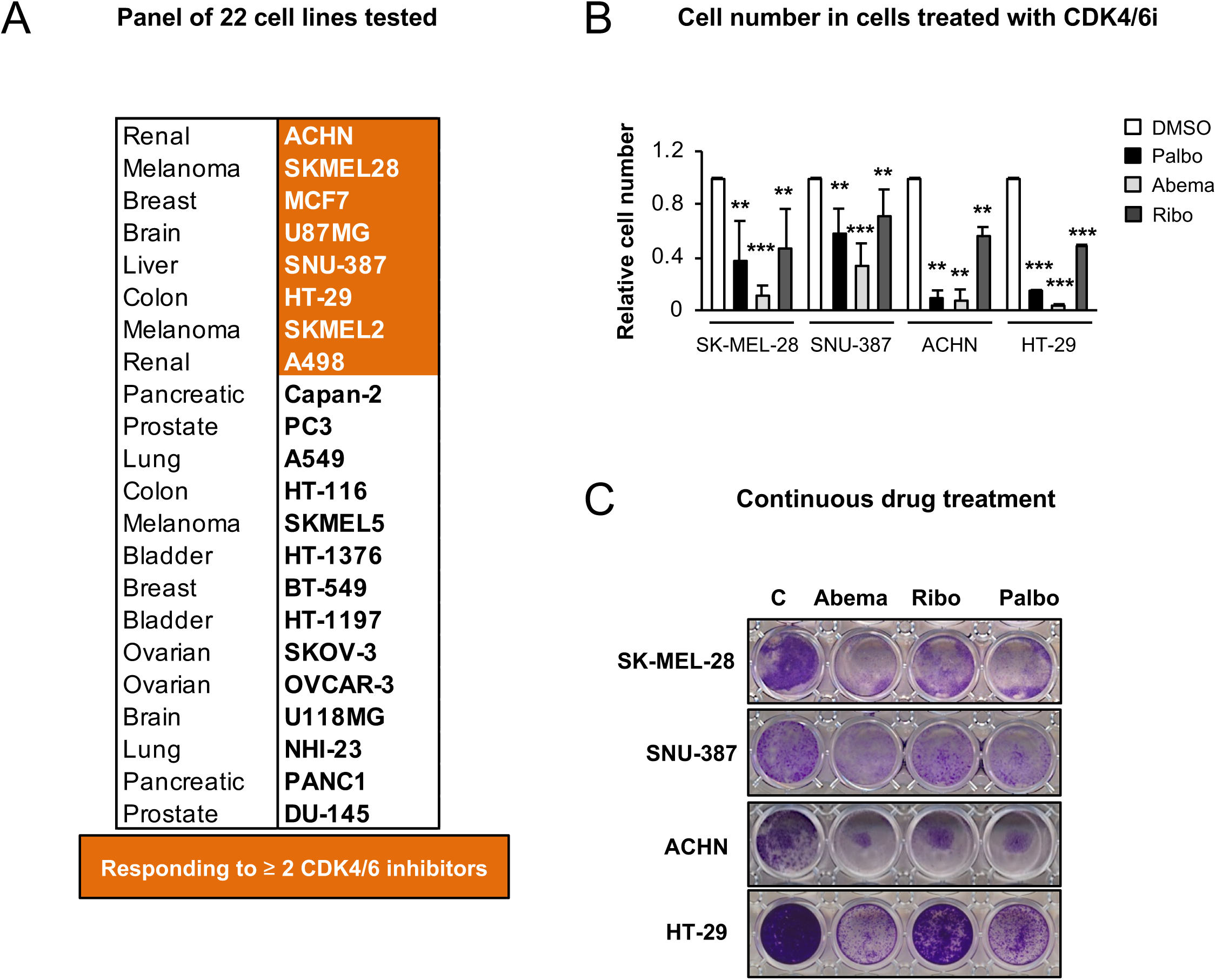
Response of other cancer cell lines to different CDK4/6 inhibitors. (**A**) Panel of cancer cell lines used in the Primary Screen to determine the efficacy of increasing concentrations of other CDK4/6/ inhibitor on proliferation. The cell lines highlighted in orange are the ones that respond to two or more inhibitors. **(B)** Quantification of relative cell number in 8 different cell lines selected for the Secondary Screen that responded to more than two CKD4/6 inhibitors. 1μM CDK4/6 inhibitor concentration was used. Data show the mean ± SD of 2-5 independent experiments. Student’s t-test analysis was performed. **(C)** Crystal violet staining showing the effect on proliferation of Abema, Ribo and Palbo in selected cell lines after 14 days of continuous drug treatment. Related to Figure 5.

## Notes

### Competing Interest Statement

The authors have declared no competing interest.

